# Unsupervised Clustering characterizes the CPG dinucleotides distribution of the Andes hantavirus

**DOI:** 10.1101/2021.02.23.432596

**Authors:** Emilio Mastriani, Shu-Lin Liu

## Abstract

Hantaviruses belong to the family of *Bunyaviridae*, and small mammals host them. Humans are infected either by inhaling virus-containing aerosols or through contact with animal droppings. Even if rodents host the pathogenic species and humans are dead-end hosts, they get accidentally infected. Andes Orthohantavirusus (ANDV) seems to be the only species with documented person-to-person transmission. Hemorrhagic fever with renal syndrome (HFRS) and hantavirus cardiopulmonary syndrome (HCPS) are both serious syndromes associated with hantavirus infections. For both syndromes, the mortality rate is near 40%. Decades of studies already highlighted the CpG repression in RNA viruses, and both the estimation of the CpG odds ratio and the correlation with their genome polarity were dominant factors to figure out the CpG bias. We conducted the differential analysis of the CpG odds ratio for all the orthohantaviruses on the full segmented genomes (L, M, S). The results suggested the statistical significance of the three groups. The *“Small”* genomes resulted in the more informative from the CpG odd ratio point of view. We calculated the CpG odds ratio for all the Orthohantaviruses within these segments, and besides, we estimated the correlation coefficient with the relative coding sequences (CDS). Preliminary results firstly confirmed the CpG odds ratio as the lowest among all the nucleotides. Second, highlighted the Andes virus as that whose CpG odds ratio within CDS is highest. The use of these two measures as features for unsupervised clustering algorithms has brought to the identification of four different sub-groups inside of the *Orthohantaviridae* family. The evidence is that the Andes Hantavirus exhibits a peculiar CpG odds ratio distribution, probably linked to its unique prerogative to pass from human-to-human.

## Introduction

Hantaviruses are enveloped RNA viruses with the negative-sense, tri-segmented genome. The large (L), the medium (M), and the small (S) segments code for viral transcriptase or polymerase, glycoprotein precursors (GPC), and the N protein that makes up the nucleocapsid, respectively [1]. Hantaviruses are transmitted to humans by infected rodents without causing any significant illness in them.

Hantavirus cardiopulmonary syndrome (HCPS) is an acute, severe, and sometimes fatal respiratory disease caused by an infection from Andes orthohantavirus. Initial symptoms are linked to the respiratory apparatus (shortness of breath, progressive cough, and tachycardia), muscle aches, fatigue, and fever, making it difficult to distinguish from simple flu. HCPS symptoms can quickly evolve and, in extreme cases, infected individuals may be incubated and receive oxygen therapy [2]. Complications of cardiogenic shock, lactic acidosis, and hemoconcentration can cause death within hours of hospitalization. In South America, Andes hantavirus (ANDV) is the primary etiologic agent. In Chile, in the period fall between 2001-2009, the authorities reported over 600 cases of ANDV-related hantavirus, with a fatality rate of 36%.

### Andes hantavirus

The Andes Orthohantavirus (ANDV) is a major causative agent of hantavirus cardiopulmonary syndrome [3]. The hantavirus cardiopulmonary syndrome is a severe respiratory disease with a fatality rate of 35–40% [4]. Andes orthohantavirus is the only hantavirus that can spread from human to human by bodily fluids or long-term contact [5–7]. The Andes virus causes the HPS into human hosts and has been identified for the first time in 1995 in samples from patients in southern Argentina [8], even if sporadic cases of HPS have been retrospectively identified [9] from as early as 1987. In 1995, for the first time, doctors identified the Andes virus in the lungs of a patient from El Bolson, and the outbreak studied in a past dispatch began on September 22, 1996. Oligoryzomys spp. rodents appear to be the principal reservoirs for most Andes viruses [10]. A previous study [11] presented the N. spinosus mice as a reservoir for the Andes virus variant found in Madre de Dios and Puno.

### CpG dinucleotides in RNA viruses

The CpG sites are regions of DNA or RNA where a guanine nucleotide follows a cytosine nucleotide in the linear sequence of bases along the direction. CpG sites occur with high frequency in genomic regions called CpG islands. CpG dinucleotides occur with a much lower frequency in the sequence of vertebrate genomes than would be expected due to random chance. This under-representation is a consequence of the high mutation rate of methylated CpG sites. The spontaneously occurring deamination of methylated cytosine results in thymine and the resulting G: T mismatched bases are often improperly resolved to A: T; whereas the deamination of cytosine results in uracil, which as a foreign base is quickly replaced by a cytosine (base excision repair mechanism). The transition rate at methylated CpG sites is folded higher than at unmethylated sites. Thus, we consider the over-representation of CpA and TpG as a consequence of the under-representation of CpG. CpG has also been observed to be predominantly under-represented in RNA viruses [12, 13] and the mechanism that contributes to the deficiency in the case of riboviruses (RNA nucleic acid) is largely unknown. Because riboviruses do not form DNA intermediates during genome replication, the methylation-deamination model is unlikely to apply, while the host innate immunity model evasion seems to be more appropriate. The CpG odds ratio values of mammals-infecting riboviruses are lower than the riboviruses infecting other taxa and the CpG motif in an AU-rich oligonucleotide can significantly stimulate the immune response of plasmacytoid dendritic cells [14]. Previous research also pointed out the huge variations of CpG bias in RNA viruses and brought out the observed under-representation of CpG in RNA viruses as not caused by the biased CpG usage in the non-coding regions but determined mainly by the coding regions [15].

This study aimed to understand whether the CpG odds ratio of Andes hantavirus is peculiar in some way. From a cluster perspective, the study also intended to verify whether the Andes hantavirus constitutes an isolated-cluster. The confirmation that this dinucleotide odds ratio is such characterizing to discriminate between groups of viruses into the same family could help to define the role of CpG islands in orthohantaviruses. Both the recurrent manifestation of the acute pulmonary syndrome in America and the urgency to understand why the Andes virus is the unique anthroponotic orthohantaviridae make research necessary.

## Materials and Methods

The genomic data to accomplish the current study have been downloaded from the ViPR [16] database. Tables 6–8 in the Appendix section report the complete list of the RNA sequences we treated. In detail, we collected 27 RNA sequences of the large genome, 39 sequences of RNA from the medium-sized genome, and 170 small genomic RNA sequences, for a total of 236 genomic segments from the Hantaviridae family. We used R version 3.6.2 and Bio Python version 1.71 to conduct the statistical analysis and calculate the CpG odds ratio, respectively. Figure 16 in the Appendix section depicts the steps followed to obtain the CpG odds ratio for all the segmented genomic sequences. Figures 17–18 in the Appendix section report the scripts used to conduct the ANOVA analysis and the unsupervised clustering in R.

### Statistical significance

Taking as null hypothesis (*H*_0_) that the means values of the CpG odds ratio from the three groups (L, M, and S) is equal, we applied for the analysis of variance (ANOVA) to accept or reject *H*_0_. To check the *normality property*, we based on the formality tests of Shapiro-Wilk with the α=0.05, while the QQ plot-chart supported our analysis as a graphical method.

Levene’s test for *homogeneity* of variance has been performed using the traditional mean-centered methodology and the R’s default median centered methodology.

### Dunn test for multiple comparisons of groups

Dunn’s Multiple Comparison Test [17, 18] is a post hoc (e.g., after an ANOVA) nonparametric test. The function we used (dunn.test) performed multiple pairwise comparisons based on Dunn’s z-test-statistic approximations to the actual rank statistics. Several options were available to adjust p-values for multiple comparisons, including methods to control the family-wise error rate (FWER) and check the false discovery rate (FDR). We used the Bonferroni adjustment (FWER) to verify Dunn’s test results and adjusted p-values = max (1, pm).

### Statistical clues to identify the most significant group concerning CpG frequency

We investigated for extra statistical clues to identify the more significant group for the CpG odds ratio. In our perspective, one group is more meaningful whether it presents a wider range of variation for the CpG odds ratio than the others.

#### Average and median of variances for the dinucleotide odd ratio

Let us introduce two measures that we will use in the coming section. Equation 1 defines the average value of the variances for the odds ratio. We calculated the value over all the dinucleotides (n=16) into each group as:

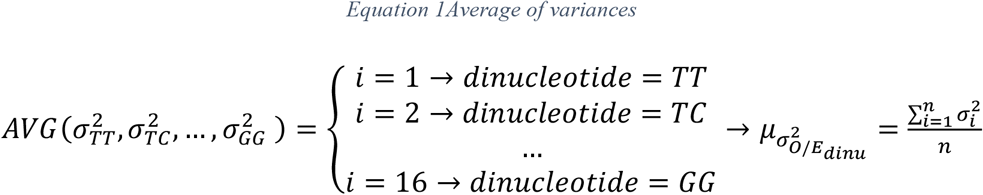

Equation 2 defines the median value along with the variances of all the odds ratios. We calculated the value over all the dinucleotides (n=16) into each group as:

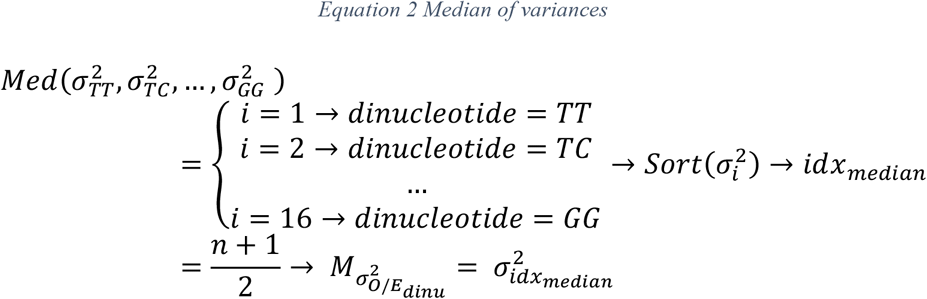

Equation 3 introduces the concept of distance between the variance of the odds ratio (calculated for the CpG dinucleotide) and the average value of all the variances of frequencies (assessed for all the dinucleotides):

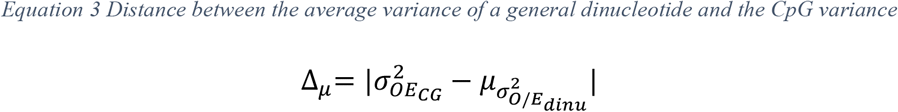

Finally, equation 4 represents the *distance* between the median of the odds ratio (calculated for the CpG dinucleotide) and the median value of all the variances of frequencies (estimated for all the dinucleotides):

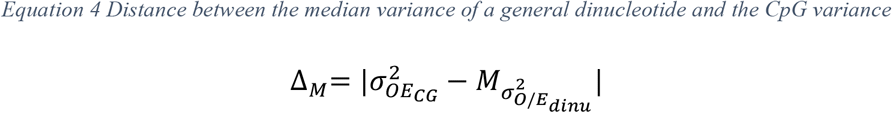

### Unsupervised clustering

To find the optimal number of clusters, we used the following four different approaches: Elbow curve method, Silhouette score method, Gap statistic method, and clustree discovery [19–21]. We executed the K-means, DBSCAN, and HCA algorithms to identify the groups of similar Hantaviruses. The CpG odds ratio, both from the full genome and from the CDS regions, and their median values from the group of small genomic segments determined the grade of similarity.

## Results

### Segmented genome and statistical difference of CpG odds ratio

#### Normality and homogeneity tests suggest to use non parametric method

Results of checking for the normality property suggested performing an equivalent non-parametric test such as a Kruskal-Wallis Test [22], not requiring the normality assumption. Table 1 reports the results for the normality test. It shows the P-value < 0.05 for the three groups, indicating that the data are not normally distributed. The QQ plots in Figure 1 reports the distribution of the CpG odds ratio for all three groups. As an assumption, the vast majority of points should follow the theoretical normal reference line and fall within the curved 95% bootstrapped confidence bands to be admitted normally distributed, and this is not the case.

**Table 1.**
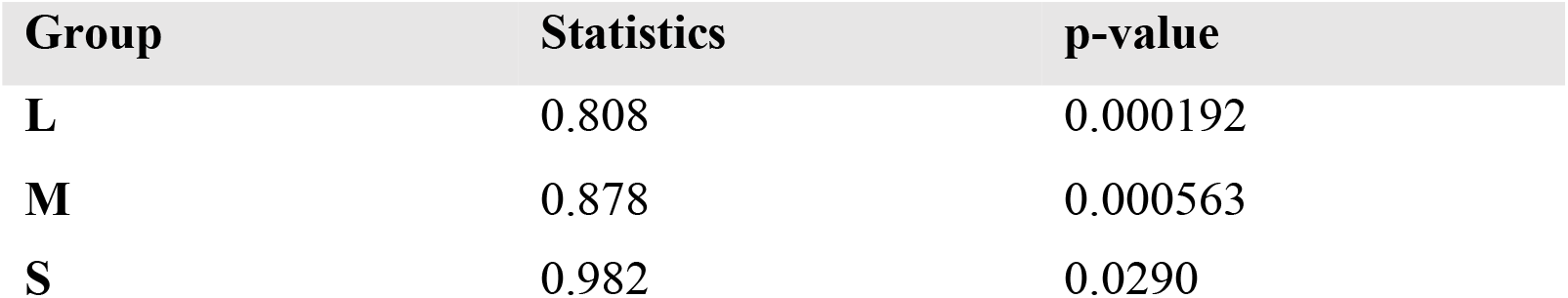
Normality test performed using Shapiro-Wilk approach for the three genomic segments type, Large (L), Medium (M), and Small (S).

**Figure 1.**
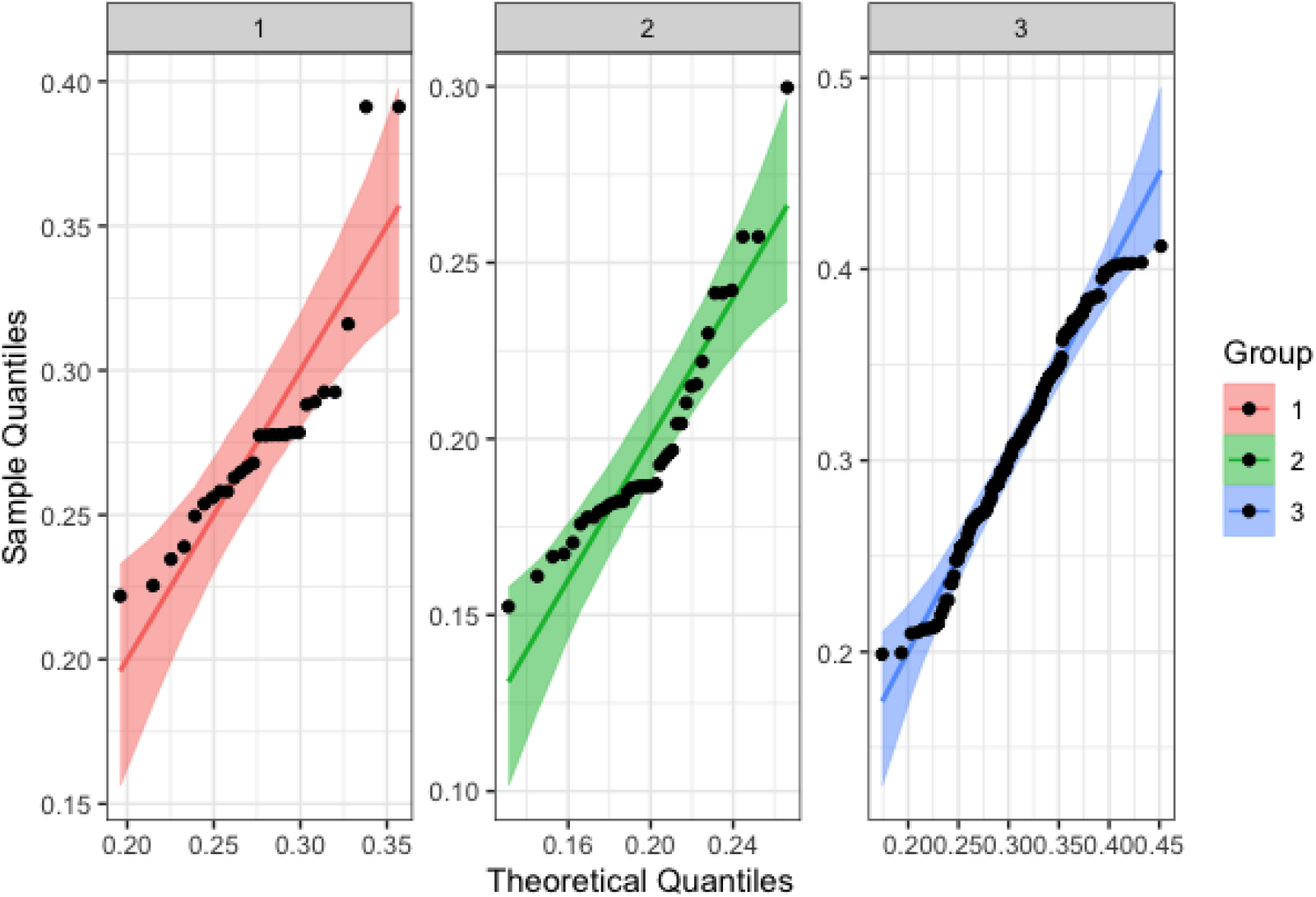
Normality QQ plots, 1 stay for group L, 2 for Medium and 3 for Small respectively

Results from Levene’s Test for Homogeneity of variance indicate that we must reject the null hypothesis and conclude that variances are not equal for at least one of our groups. Table 2 displays the test statistic for two different versions of Levene’s test. A p-value = 0.0003128 or 0.0001199 tends to accept the alternative hypothesis of inequality variances. The boxplot reported in Figure 2 also indicates some major outliers, enough evidence to use the Kruskal-Wallis ANOVA as a non-parametric test.

**Table 2.** Test of homogeneity of variance

| *Levene's Test for Homogeneity of Variance (center = "mean")* |  |  |
| --- | --- | --- |
| *Df* | F value | Pr (>F) |
| 2 | 8.356 | 0.0003128 |
| *Levene's Test for Homogeneity of Variance (center = "median")* |  |  |
| 2 | 9.3875 | 0.0001199 |

**Figure 2.**
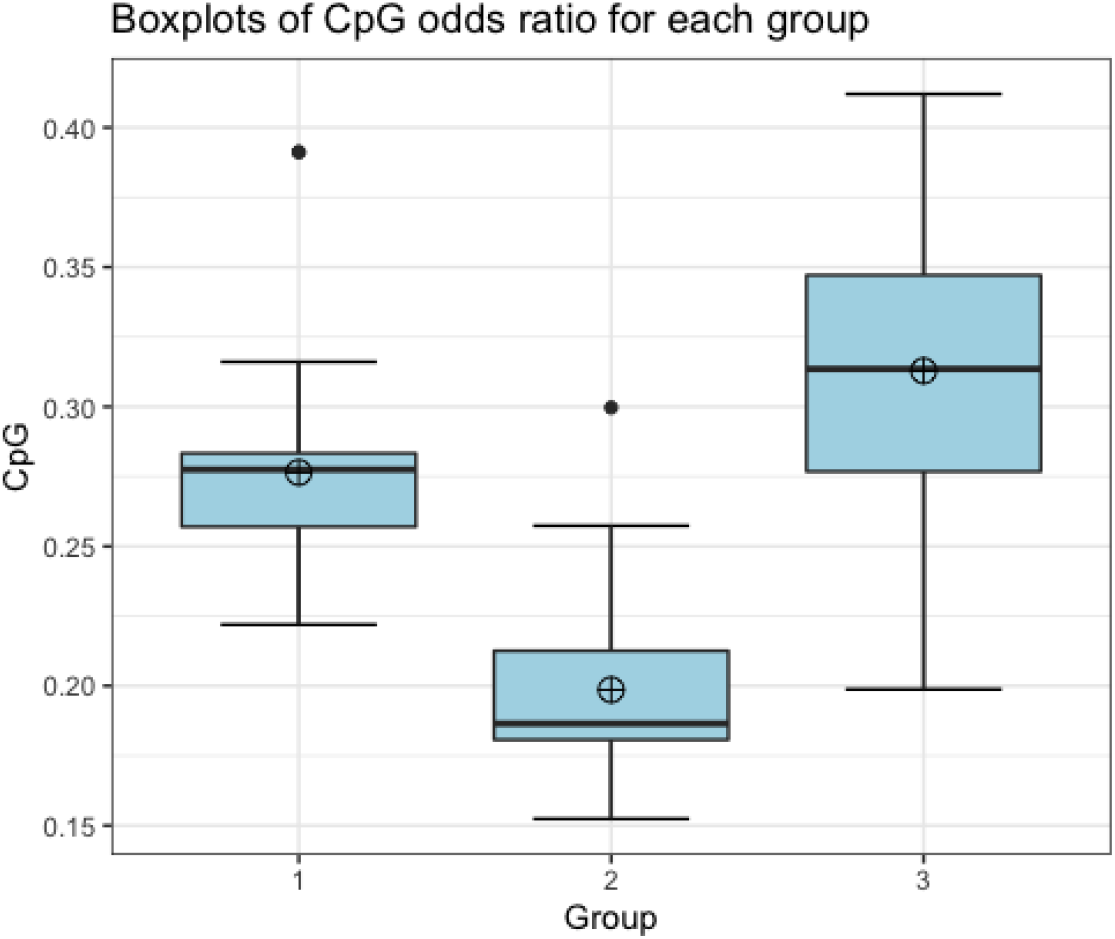
Boxplots to Visually Check for Outliers. 1 stay for group L, 2 for Medium and 3 for Small respectively

**Figure 3.**
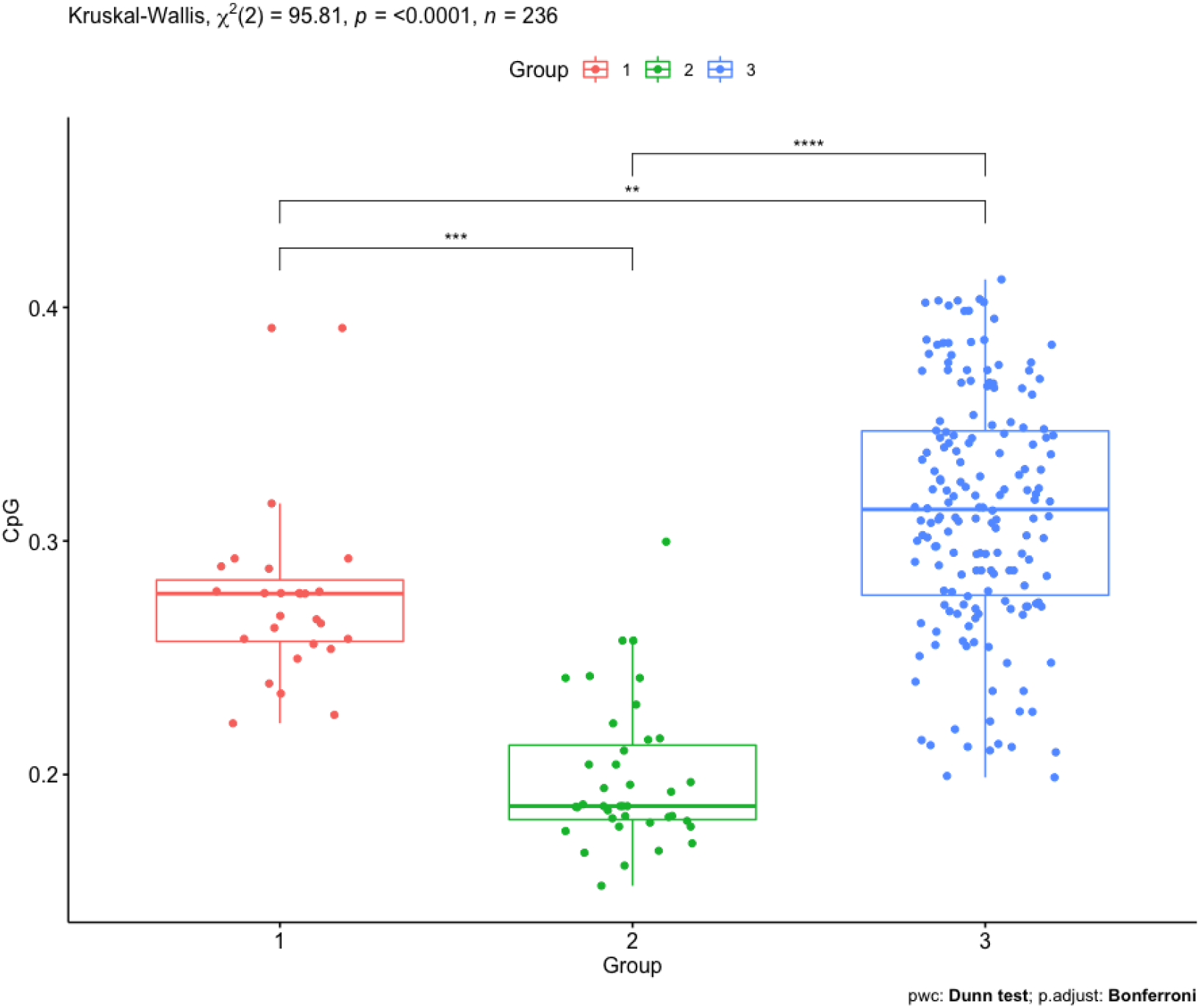
Boxplots representation of the Dunn’s test

**Figure 4.**
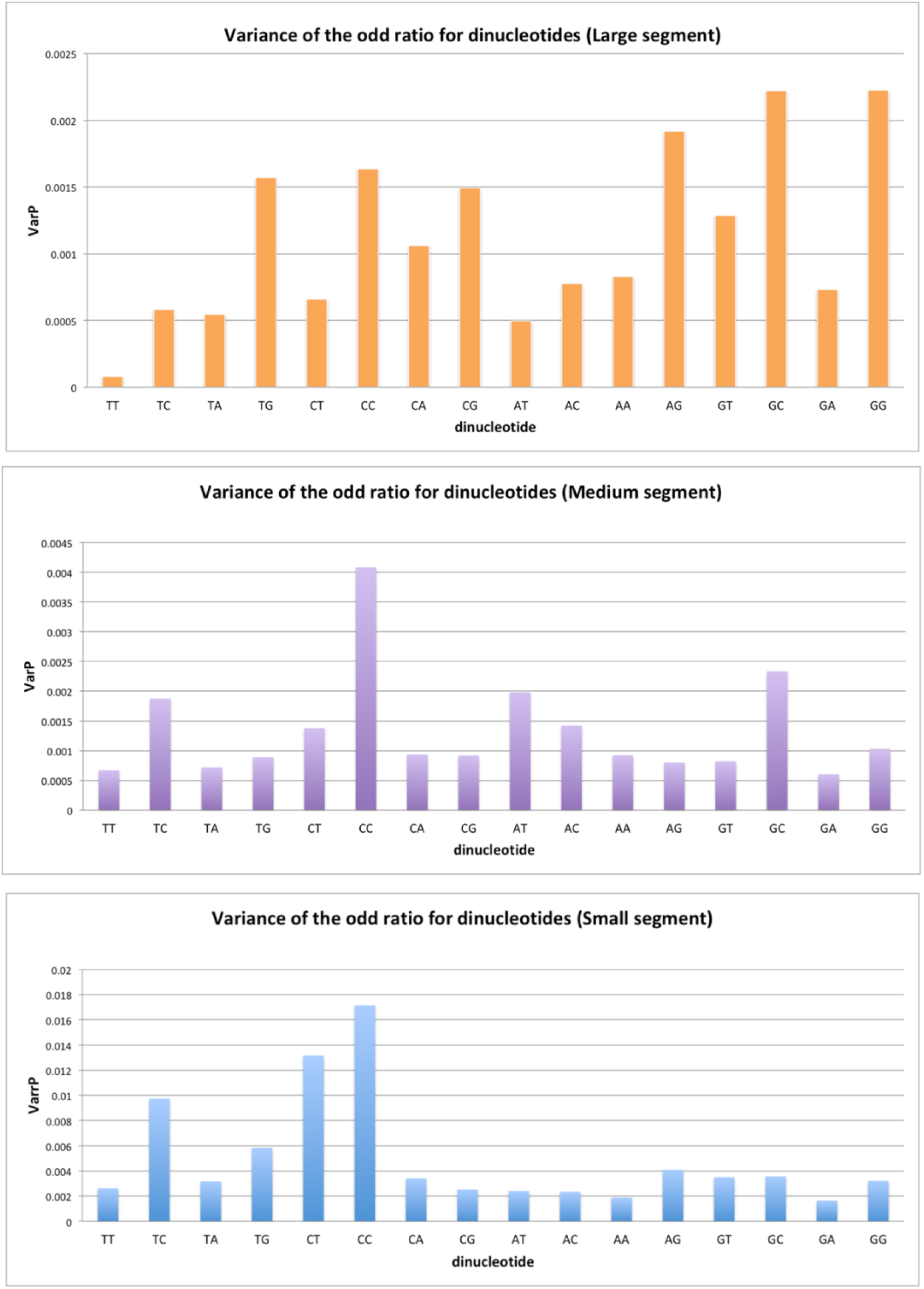
Variance of the dinucleotide frequency for the three genomic groups (L, M and S)

#### Kruskal-Wallis test suggests to reject the null hypothesis

The Kruskal-Wallis test results in a two-sided test *p – value* < 2.2e^−16^. This indicates that we should reject the null hypothesis that means ranks are equal across groups and conclude that there is a significant difference in CpG odds ratio distribution. Descriptive statistics indicate that the median value with 95% confidence intervals for group L is 0.277, group M is 0.187, and group S is 0.314. That is to say, the difference between the median values of each segment L and M is about 0.09 (p=1.137969e-04), segments L and S are about 0.037 (p=7.471942e-04), and segments M and S is about 0.127 (p=2.173163e-21).

### The small genomics segments clique as the more informative group

#### Dunn test result: significant difference in the CpG odds ratio between the L, M, and S groups

Dunn’s Multiple Comparison method tests stochastic dominance and reports the results among multiple pairwise comparisons after a Kruskal-Wallis test among k groups. The null hypothesis for each pairwise comparison is that the probability of observing a randomly selected value from the first group *is* larger than a randomly selected value from the second group equals one-half, and so rejecting *H*_0_ based on p≤α/2. Table 3 reports the result from the Dunn’s test providing all possible pairwise comparisons. In the table, the adjusted *p*-values will have an asterisk next to them we would reject the null hypotheses at the specified significance level, comparisons rejected with the Bonferroni adjustment at the α level (two-sided test). Figure 7 shows the test output between groups. If we consider the total distance from each group to the others, then it suggests that the difference between group n. 3 (Small segments) and the other groups is more significant.

**Table 3.**
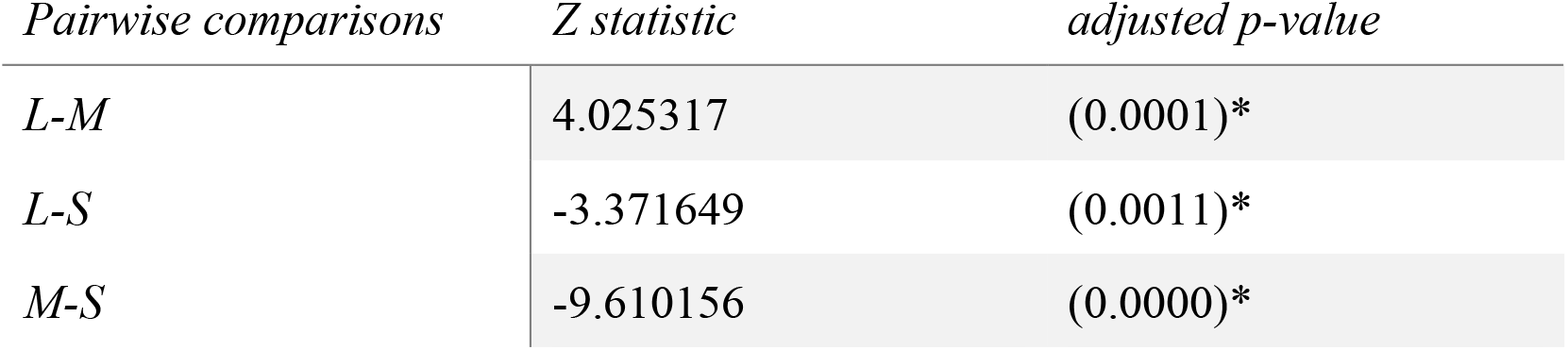
Kruskal-Wallis rank sum test. Comparison of x by group

| Pairwise comparisons | Z statistic | adjusted p-value |
| --- | --- | --- |
| L-M | 4.025317 | (0.0001)* |
| L-S | -3.371649 | (0.0011)* |
| M-S | -9.610156 | (0.0000)* |

#### The variance of the CpG dinucleotide’s odd ratio in the group of small genomic segments will be biologically relevant

In our minds, one group could be more meaningful than one other whether it presents a wider range of variation for the CpG odds ratio. Variance (σ^2^) is a measurement of the spread between numbers in a data set. It measures how far each number in the set is from the mean and consequently from every other number in the collection. Figure 8 reports the variance of the odds ratio for every dinucleotide in each group of genomic segments. Results show as the CpG di-nucleotide tends to be more conservative in comparison with the other di-nucleotides. Also, the group of small genomic segments, with a value close to 0.002, presents the lowest CpG odds ratio variation. The outcomes suggest that possible variation inside of this group should be biologically relevant.

#### The group of small genomic segments is the more informative from the CpG odds ratio point of view

Figure 5 compares the odds ratio variance of CpG dinucleotide, the average and median variance for any dinucleotide, grouped by genomic segments (L, M, and S). While the measures do not represent meaningful differences for large and medium genomic segments, the small genomic group depicts a more unusual situation. The value of the variance for the small genomic CpG is far away from the median and average values of the dinucleotides from the other groups. The observation becomes more evident from the diagram in Figure 6, indicating the *distance* values. For each group of genomic segments (L, M, and S), we estimated the following measures:

1. 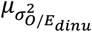, the average of the variance for all the dinucleotides
2. Δ_*μ*_, the distance of CpG odds ratio variance from 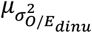
3. Δ_*M*_, the distance of CpG odds ratio variance from 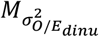

**Figure 5.**
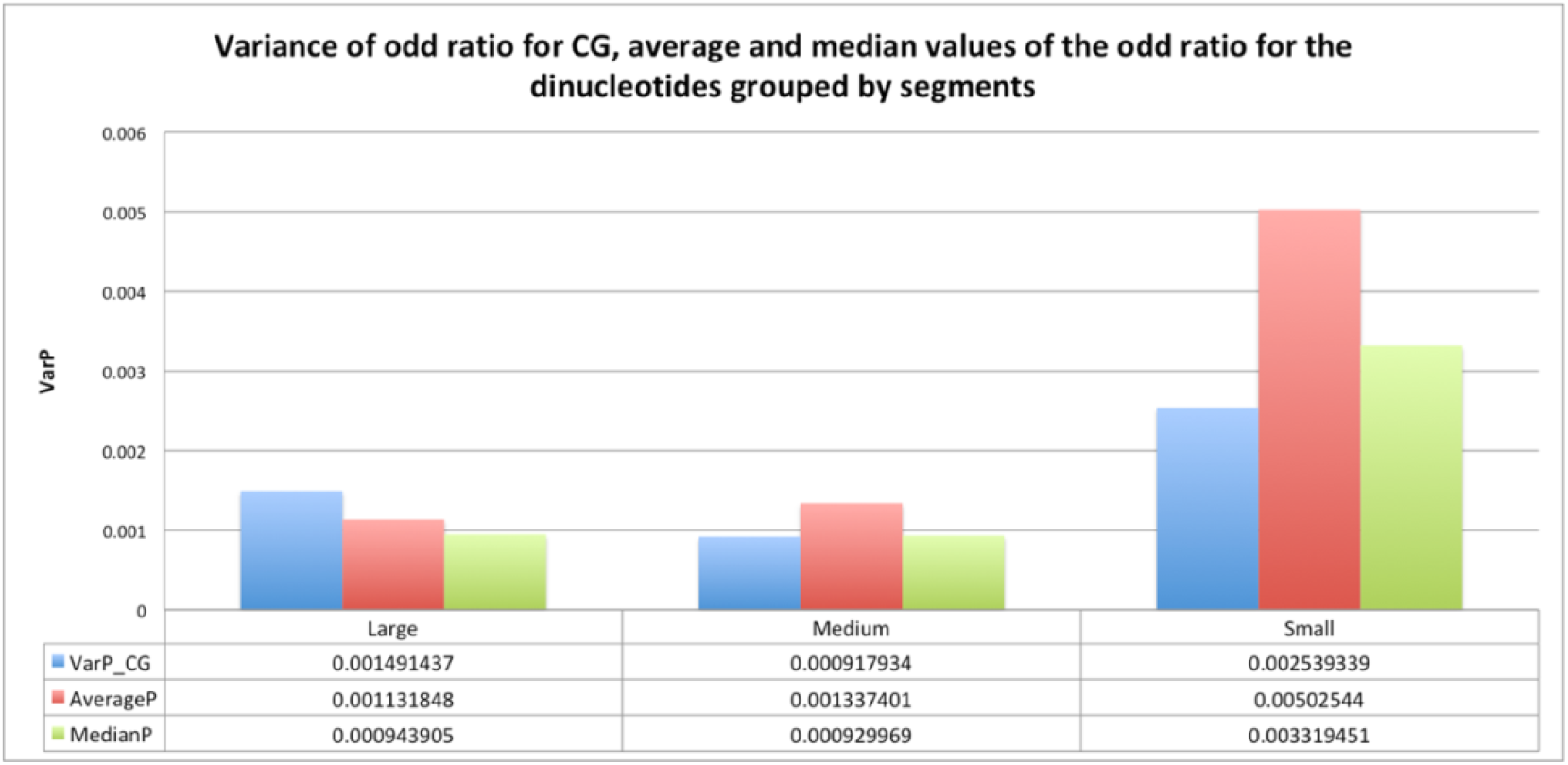
Comparison between the odds ratio variance of CpG dinucleotide and the average and median variance for generic dinucleotide grouped by genomic segments (L, M and S).

**Figure 6.**
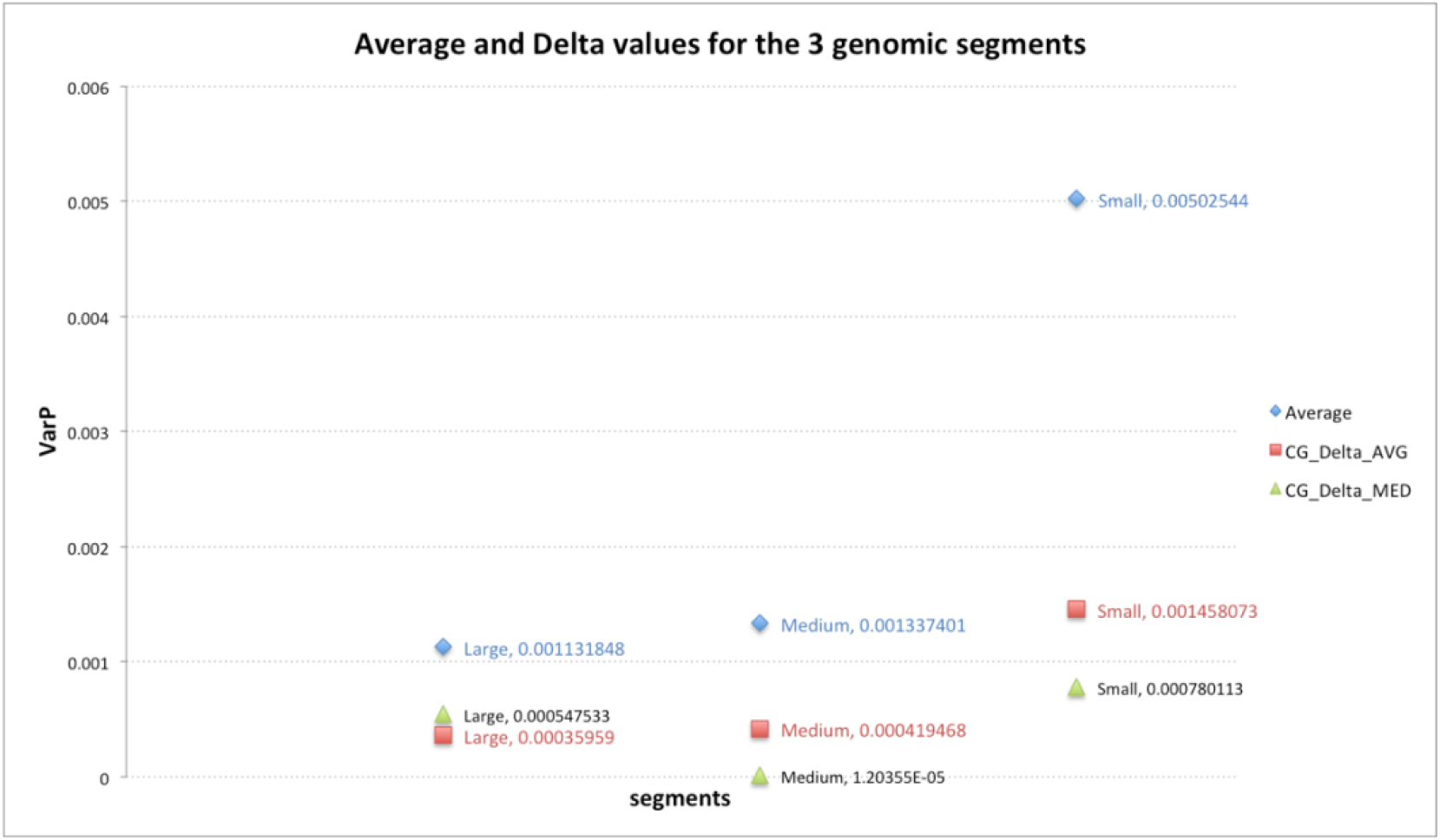
Comparison of the distances between the average of the variance for all the dinucleotides (Average, blue diamond), the distance of CpG odds ratio variance from the Average measure (CG_Delta_AVG, red square) and the distance of CpG odds ratio variance for all the dinucleotides (CG_Delta_MED, green triangle). The vales are grouped by genomic segment type (L, M and S)

The consideration of those measures for the three groups (L, M, and S) drives to the following observations: 1. a higher value than the average value of the variances of the odds ratio over all the di-nucleotides compared to the other groups indicates that within the small group the odds ratio tends to change more frequently; 2. a higher value than the distance between the variance of the odds ratio for the CpG dinucleotide and the average value of all the variances of all the frequencies for all the di-nucleotides compared to other groups shows that within the small group CpG hold the highest rate of variability; 3. a higher value than the distance between the median of the odds ratio for the CpG dinucleotide and the median value of all the variances of all the frequencies for all the di-nucleotides compared to the other groups shows that within the small group even the CpG median has the most representative value. These results indicate that the group of small genomic segments is the more informative from the CpG odds ratio point of view.

### Influence of the CpG odds ratio from regions

#### The CpG Odds ratio inside CDS regions of short genomic segments is the lowest for the Hantaviridae family

Previous studies already underlined the CpG odds ratio as the lowest compared to those of the other dinucleotides, even in the case of RNA viruses [15]. The calculation of the odds ratio for all the dinucleotides around the CDS regions restricted our study to ten different RNA viruses from the *Hantaviridae* family: Andes, Tunari, Bayou, Choclo, Dobrava-Belgrade, Hantaan, Hantaanvirus, Puumala, Seoul, and Tula. This computation confirmed the CpG odds ratio into CDS as the lowest also for a group of small genomic segments, as reported by Figure 7. It shows that the odds ratio for CpG in CDS regions is the lowest compared to the odds ratio of other dinucleotides for the ten considered viruses.

**Figure 7.**
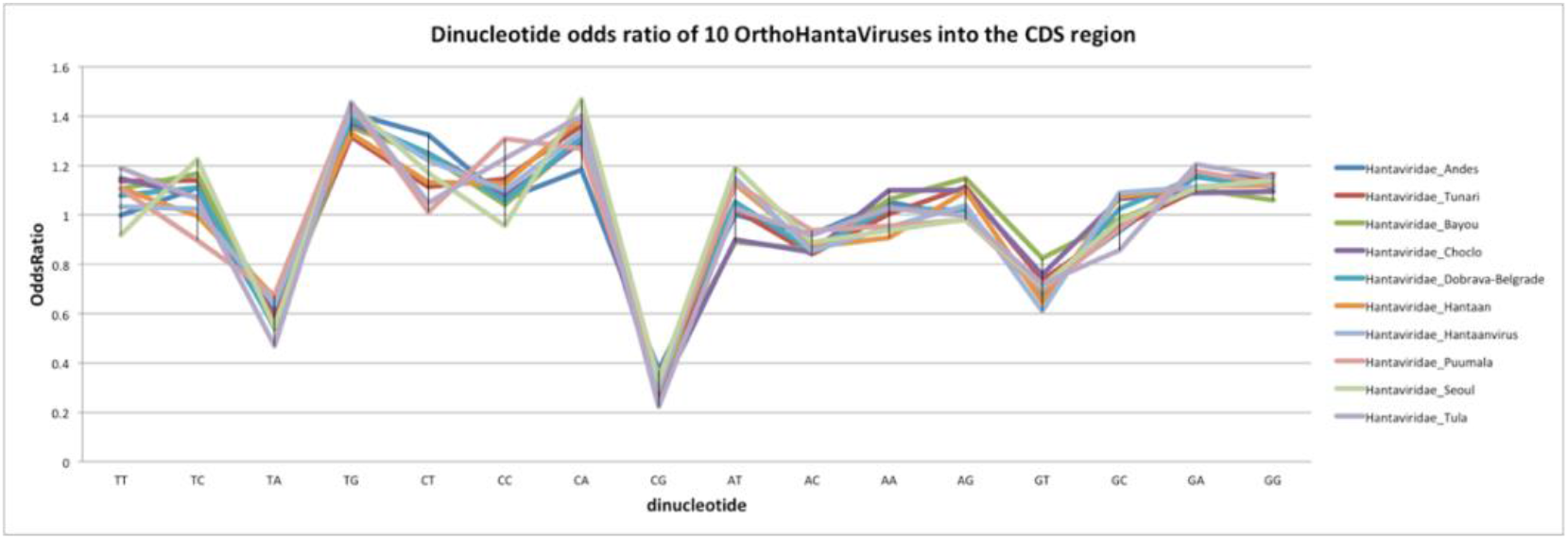
Dinucleotide odds ratio into CDS regions for the 10 viruses. The CDS regions belong to the group of small genomic segments

#### Andes hantavirus and CpG frequency from CDS regions

Dealing with the frequency of CpG inside the coding regions, make evident how *Hantaviridae* Andes can suggest itself as a specific case. We calculated the Pearson correlation coefficient from the data reported in Table 4, getting a rate near 0.98. Such a result supports the positive correlation between the CpG odds ratio over the full genome and the CpG odds ratio in the CDS. This result highlights that the CpG dinucleotides perform a function in the coding regions. The data collected in Table 4 shows how the CpG frequency in CDS for the Andes *Hantaviridae* represents the highest value, 7.58% greater than the second-highest value (*Hantaviridae* Dobrava-Belgrade). The odds ratio bars depicted in Figure 8 prove that the Andes H. compared with the other hantaviruses, detains the highest CpG odds ratio into CDS regions.

**Table 4.**
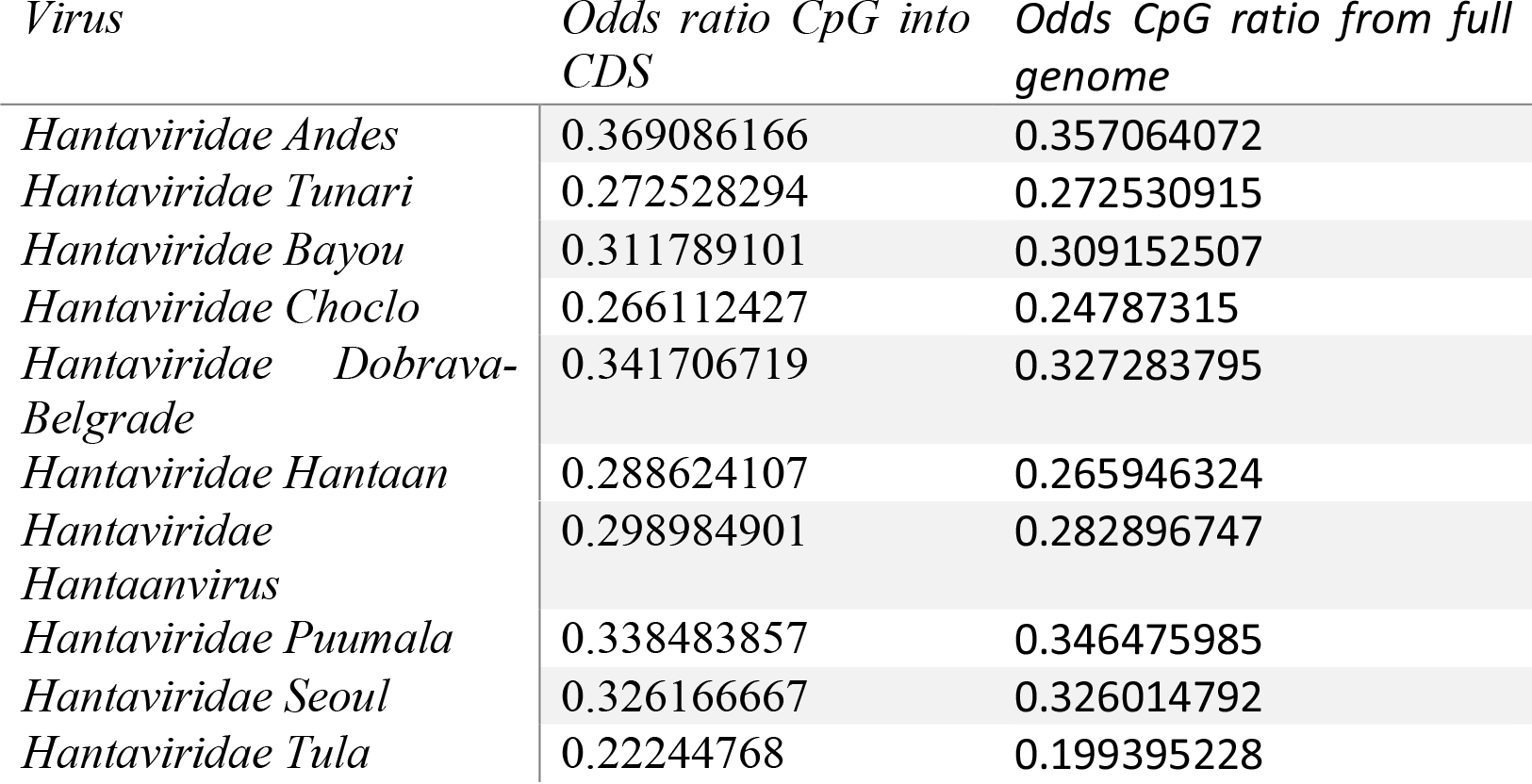
CpG odds ratio from CDS regions and from full genome into the group of small genomic segments

**Figure 8.**
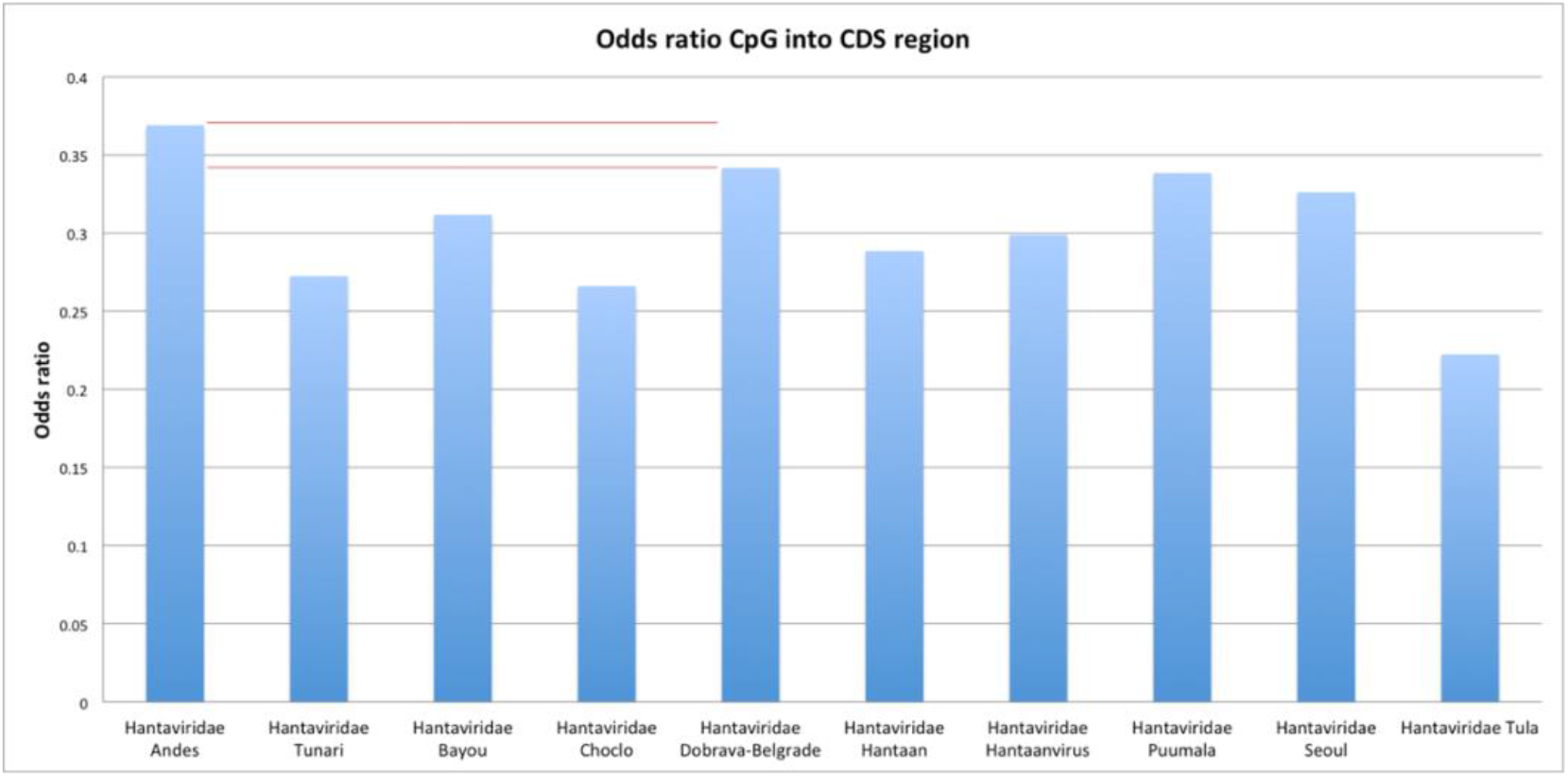
Odds ratio of CpG into CDS regions

Examining the odds ratio of CpG along the full genome, the CpG odds ratio into the CDS regions, and the median value for hantaviruses pointed out above, the Hantaviridae Andes holds on the highest rates for all three measures. Table 5 and Figure 9 show that the CpG odds ratio value into CDS of *Hantaviridae* Andes is 7.58% greater than the same value from *Hantaviridae* Dobrava-Belgrade (the second virus sorted by CpG odds ratio into CDS value). And again, *Hantaviridae* Andes is 3.08% and 5.78% greater than *Hantaviridae* Puumala (the second virus for CpG into the full genome and CpG median values), respect to the CpG into the full genome and CpG median values. In conclusion, the Andes *Hantaviridae* brings the highest value of CpG odds ratio into CDS regions. The Pearson correlation close to 0.98 confirms that the CpG odds ratio along the full small genomic segment and the CpG odds ratio into the CDS regions of the same genomic segment are positively related. This clue stresses the possible roles carried out by the CpG islands into the coding regions. Lastly, by facing the CpG odds ratio from the full genome, from the CDS regions, and the median values, draw attention to a stronger concentration of CpG islands both along the full small genomic segment and into the CDS regions for the Andes virus.

**Table 5.** Comparison (Δ) of the CpG odds ratio in CDS, CpG odds ratio from full genome and Median values for the viruses with the top frequencies

| | Andes | Dobrava-Belgrade | Puumala | $\Delta$ |
| --- | --- | --- | --- | --- |
| CpG into CDS | 0.369 | 0.341 |  | 0.028 |
| CpG full genome | 0.357 |  | 0.346 | 0.011 |
| Median | 0.363 |  | 0.342 | 0.021 |

**Table 6.**
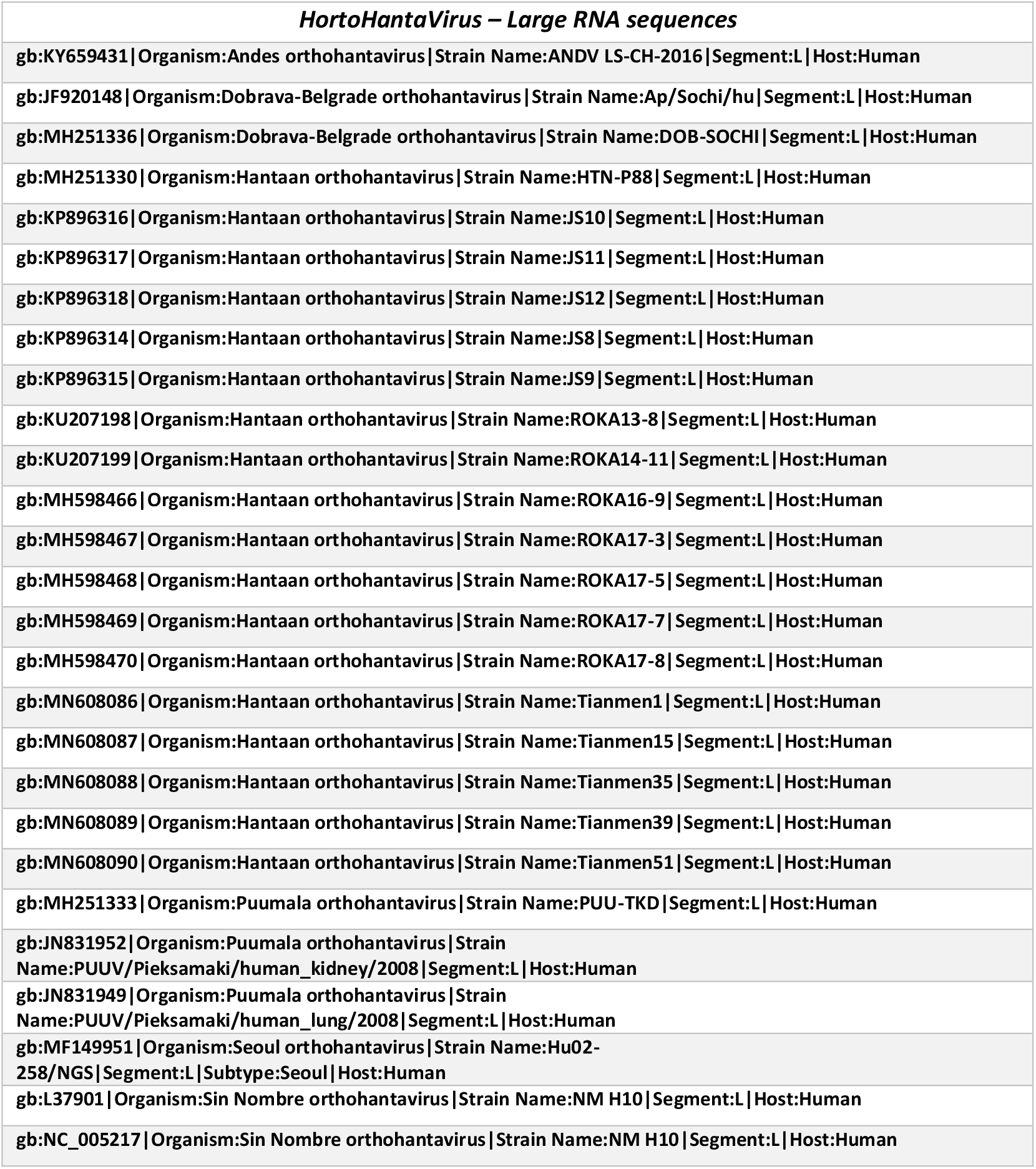
List of large RNA sequences

**Table 7.**
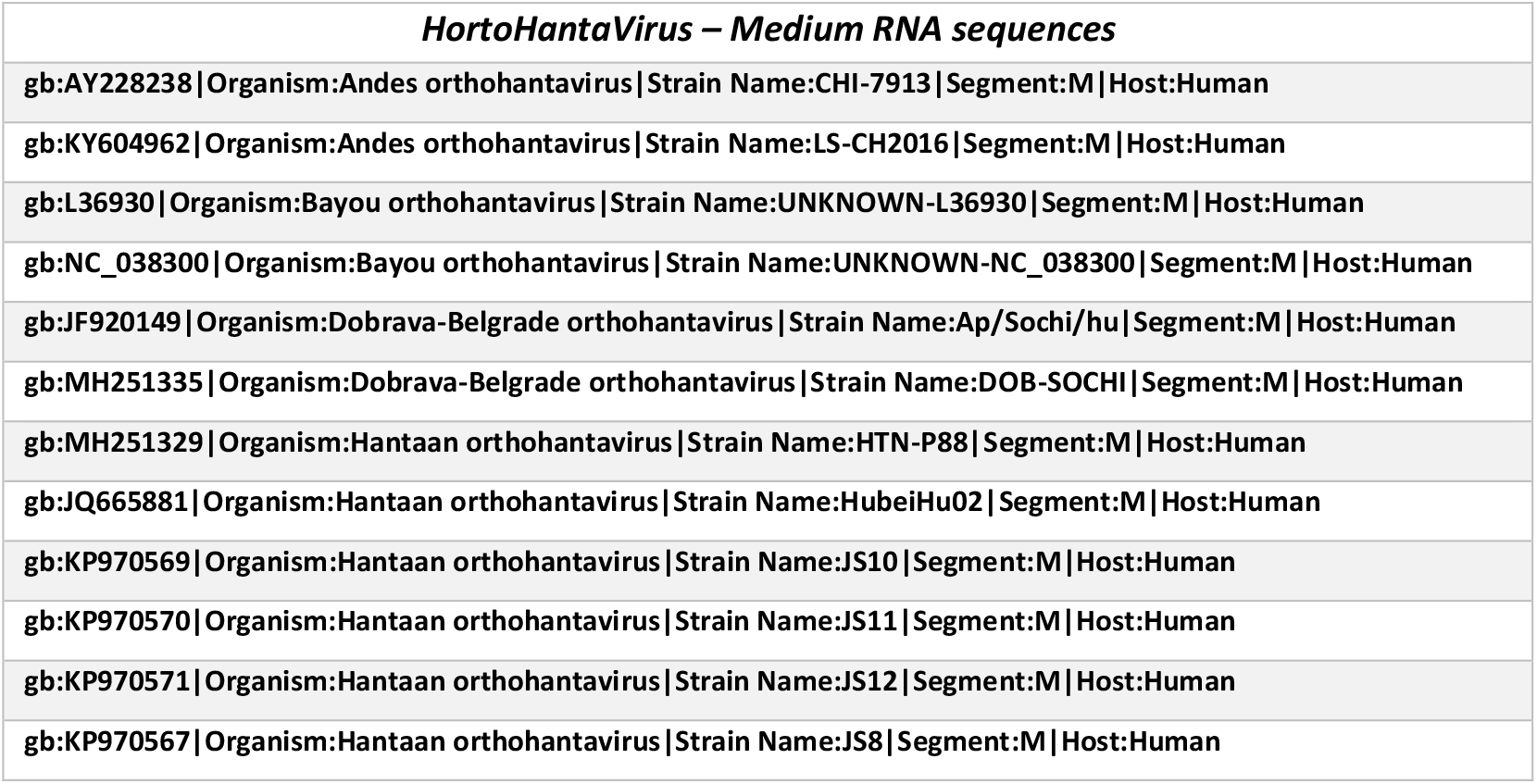
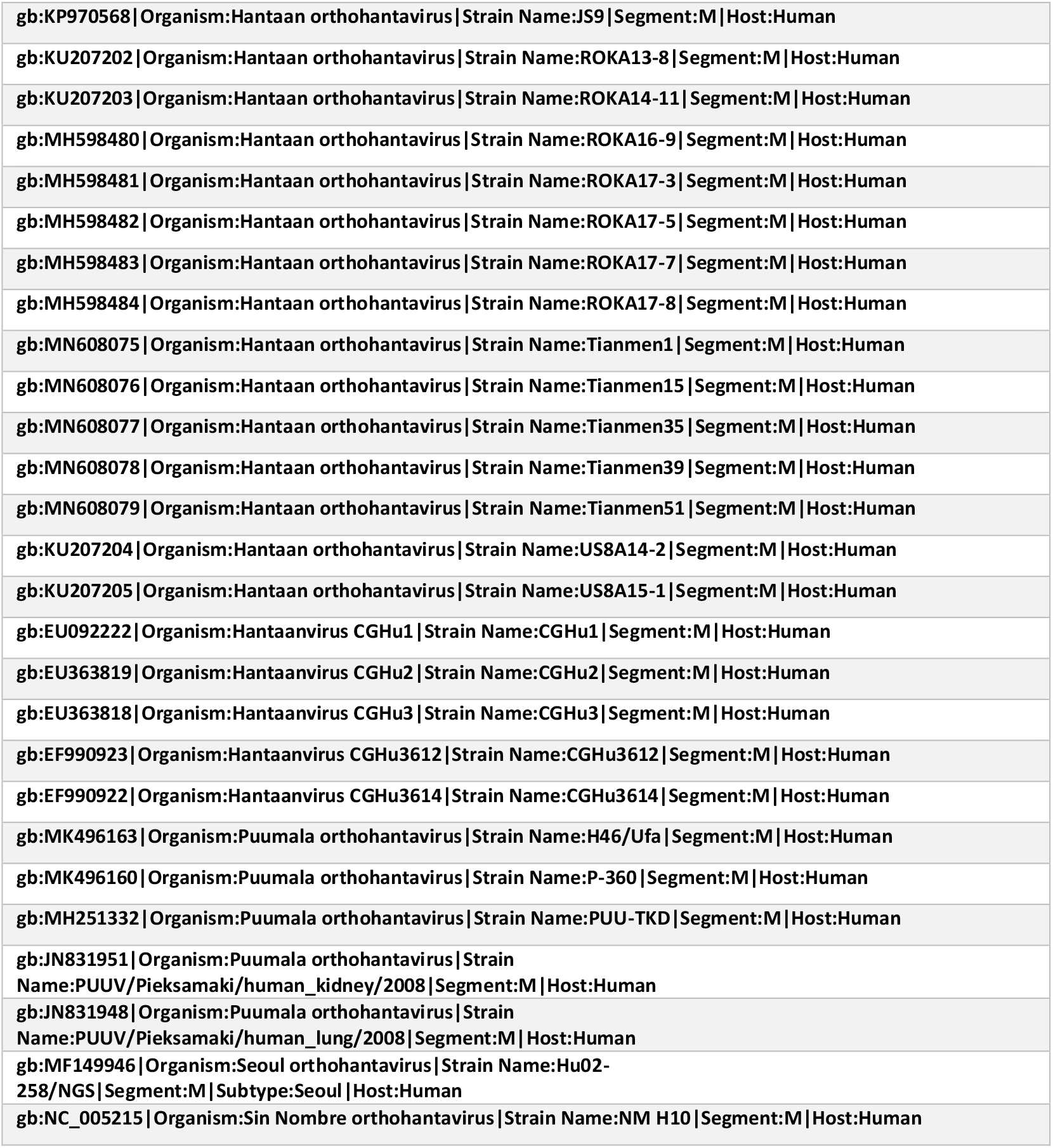
List of medium RNA sequences

**Table 8.** List of small RNA sequences

| *HortoHantaVirus – Small RNA sequences* |
| --- |
| gb:KY659432\|Organism:Andes orthohantavirus\|Strain Name:ANDV LS-CH-2016 ex Chile\|Segment:S\|Host:Human |
| gb:AY228237\|Organism:Andes orthohantavirus\|Strain Name:CHI-7913\|Segment:S\|Host:Human |
| gb:JF750419\|Organism:Tunari virus\|Strain Name:FVB554\|Segment:S\|Host:Human |
| gb:JF750418\|Organism:Tunari virus\|Strain Name:FVB640\|Segment:S\|Host:Human |
| gb:JF750417\|Organism:Tunari virus\|Strain Name:FVB799\|Segment:S\|Host:Human |
| gb:L36929\|Organism:Bayou orthohantavirus\|Strain Name:UNKNOWN-L36929\|Segment:S\|Host:Human |
| gb:NC_038298\|Organism:Bayou orthohantavirus\|Strain Name:UNKNOWN-NC_038298\|Segment:S\|Host:Human |
| gb:KM597161\|Organism:Choclo virus\|Strain Name:Uk (ex Panama)\|Segment:S\|Host:Human |
| gb:KP878313\|Organism:Dobrava-Belgrade orthohantavirus\|Strain Name:10752/hu\|Segment:S\|Host:Human |
| gb:JF920150\|Organism:Dobrava-Belgrade orthohantavirus\|Strain Name:Ap/Sochi/hu\|Segment:S\|Host:Human |
| gb:MH251334\|Organism:Dobrava-Belgrade orthohantavirus\|Strain Name:DOB-SOCHI\|Segment:S\|Host:Human |
| gb:KC570384\|Organism:Hantaan orthohantavirus\|Strain Name:DandongHu-22\|Segment:S\|Host:Human |
| gb:KC570385\|Organism:Hantaan orthohantavirus\|Strain Name:DandongHu-28\|Segment:S\|Host:Human |
| gb:KC570386\|Organism:Hantaan orthohantavirus\|Strain Name:DandongHu-32\|Segment:S\|Host:Human |
| gb:KC570387\|Organism:Hantaan orthohantavirus\|Strain Name:DandongHu-34\|Segment:S\|Host:Human |
| gb:KC570388\|Organism:Hantaan orthohantavirus\|Strain Name:DandongHu-44\|Segment:S\|Host:Human |
| gb:KC570389\|Organism:Hantaan orthohantavirus\|Strain Name:DandongHu-89\|Segment:S\|Host:Human |
| gb:KC570390\|Organism:Hantaan orthohantavirus\|Strain Name:DandongHu-91\|Segment:S\|Host:Human |
| gb:MH251328\|Organism:Hantaan orthohantavirus\|Strain Name:HTN-P88\|Segment:S\|Host:Human |
| gb:MN478382\|Organism:Hantaan orthohantavirus\|Strain Name:HTNV-HN2017/70\|Segment:S\|Host:Human |
| gb:MN478383\|Organism:Hantaan orthohantavirus\|Strain Name:HTNV-HN2017/76\|Segment:S\|Host:Human |
| gb:MN478384\|Organism:Hantaan orthohantavirus\|Strain Name:HTNV-HN2017/79\|Segment:S\|Host:Human |
| gb:MN478385\|Organism:Hantaan orthohantavirus\|Strain Name:HTNV-HN2017/80\|Segment:S\|Host:Human |
| gb:MN478386\|Organism:Hantaan orthohantavirus\|Strain Name:HTNV-HN2017/81\|Segment:S\|Host:Human |
| gb:MN478387\|Organism:Hantaan orthohantavirus\|Strain Name:HTNV-HN2017/82\|Segment:S\|Host:Human |
| gb:MN478388\|Organism:Hantaan orthohantavirus\|Strain Name:HTNV-HN2017/87\|Segment:S\|Host:Human |
| gb:MN478389\|Organism:Hantaan orthohantavirus\|Strain Name:HTNV-HN2018/106\|Segment:S\|Host:Human |
| gb:MN478390\|Organism:Hantaan orthohantavirus\|Strain Name:HTNV-HN2018/131\|Segment:S\|Host:Human |
| gb:MN478391\|Organism:Hantaan orthohantavirus\|Strain Name:HTNV-HN2018/134\|Segment:S\|Host:Human |
| gb:MN478392\|Organism:Hantaan orthohantavirus\|Strain Name:HTNV-HN2018/138\|Segment:S\|Host:Human |
| gb:MN478393\|Organism:Hantaan orthohantavirus\|Strain Name:HTNV-HN2018/146\|Segment:S\|Host:Human |
| gb:MN478394\|Organism:Hantaan orthohantavirus\|Strain Name:HTNV-HN2018/150\|Segment:S\|Host:Human |
| gb:MN478395\|Organism:Hantaan orthohantavirus\|Strain Name:HTNV-HN2018/152\|Segment:S\|Host:Human |
| gb:MN478396\|Organism:Hantaan orthohantavirus\|Strain Name:HTNV-HN2018/154\|Segment:S\|Host:Human |
| gb:MN478397\|Organism:Hantaan orthohantavirus\|Strain Name:HTNV-HN2018/157\|Segment:S\|Host:Human |
| gb:JQ665905\|Organism:Hantaan orthohantavirus\|Strain Name:HubeiHu02\|Segment:S\|Host:Human |
| gb:KP970581\|Organism:Hantaan orthohantavirus\|Strain Name:JS10\|Segment:S\|Host:Human |
| gb:KP970582\|Organism:Hantaan orthohantavirus\|Strain Name:JS11\|Segment:S\|Host:Human |
| gb:KP970583\|Organism:Hantaan orthohantavirus\|Strain Name:JS12\|Segment:S\|Host:Human |
| gb:KP970579\|Organism:Hantaan orthohantavirus\|Strain Name:JS8\|Segment:S\|Host:Human |
| gb:KP970580\|Organism:Hantaan orthohantavirus\|Strain Name:JS9\|Segment:S\|Host:Human |
| gb:KY283955\|Organism:Hantaan orthohantavirus\|Strain Name:MN2009P-M3\|Segment:S\|Host:Human |
| gb:KY283956\|Organism:Hantaan orthohantavirus\|Strain Name:MN2009P-M6\|Segment:S\|Host:Human |
| gb:KU207206\|Organism:Hantaan orthohantavirus\|Strain Name:ROKA13-8\|Segment:S\|Host:Human |
| gb:KU207207\|Organism:Hantaan orthohantavirus\|Strain Name:ROKA14-11\|Segment:S\|Host:Human |
| gb:MH598494\|Organism:Hantaan orthohantavirus\|Strain Name:ROKA16-9\|Segment:S\|Host:Human |
| gb:MH598495\|Organism:Hantaan orthohantavirus\|Strain Name:ROKA17-3\|Segment:S\|Host:Human |
| gb:MH598496\|Organism:Hantaan orthohantavirus\|Strain Name:ROKA17-5\|Segment:S\|Host:Human |
| gb:MH598497\|Organism:Hantaan orthohantavirus\|Strain Name:ROKA17-7\|Segment:S\|Host:Human |
| gb:MH598498\|Organism:Hantaan orthohantavirus\|Strain Name:ROKA17-8\|Segment:S\|Host:Human |
| gb:KC844226\|Organism:Hantaan orthohantavirus\|Strain Name:SXHu2012B1\|Segment:S\|Host:Human |
| gb:KC844227\|Organism:Hantaan orthohantavirus\|Strain Name:SXHu2012B3\|Segment:S\|Host:Human |
| gb:MN608064\|Organism:Hantaan orthohantavirus\|Strain Name:Tianmen1\|Segment:S\|Host:Human |
| gb:MN608065\|Organism:Hantaan orthohantavirus\|Strain Name:Tianmen15\|Segment:S\|Host:Human |
| gb:MN608066\|Organism:Hantaan orthohantavirus\|Strain Name:Tianmen35\|Segment:S\|Host:Human |
| gb:MN608067\|Organism:Hantaan orthohantavirus\|Strain Name:Tianmen39\|Segment:S\|Host:Human |
| gb:MN608068\|Organism:Hantaan orthohantavirus\|Strain Name:Tianmen51\|Segment:S\|Host:Human |
| gb:KU207208\|Organism:Hantaan orthohantavirus\|Strain Name:US8A14-2\|Segment:S\|Host:Human |
| gb:KU207209\|Organism:Hantaan orthohantavirus\|Strain Name:US8A15-1\|Segment:S\|Host:Human |
| gb:KM355414\|Organism:Hantaan orthohantavirus\|Strain Name:WCL\|Segment:S\|Host:Human |
| gb:KY357324\|Organism:Hantaan orthohantavirus\|Strain Name:XA2009P-M18\|Segment:S\|Host:Human |
| gb:KY357325\|Organism:Hantaan orthohantavirus\|Strain Name:XA2011P-Z21\|Segment:S\|Host:Human |
| gb:KY357323\|Organism:Hantaan orthohantavirus\|Strain Name:XA2012P-Z22\|Segment:S\|Host:Human |
| gb:KY357326\|Organism:Hantaan orthohantavirus\|Strain Name:XA2012P133\|Segment:S\|Host:Human |
| gb:KY357327\|Organism:Hantaan orthohantavirus\|Strain Name:XA2012P148\|Segment:S\|Host:Human |
| gb:KY357322\|Organism:Hantaan orthohantavirus\|Strain Name:XA2012P160\|Segment:S\|Host:Human |
| gb:HQ834507\|Organism:Hantaan virus P09072\|Strain Name:P09072\|Segment:S\|Host:Human |
| gb:EU092218\|Organism:Hantaanvirus CGHu1\|Strain Name:CGHu1\|Segment:S\|Host:Human |
| gb:EU363813\|Organism:Hantaanvirus CGHu2\|Strain Name:CGHu2\|Segment:S\|Host:Human |
| gb:EU363809\|Organism:Hantaanvirus CGHu3\|Strain Name:CGHu3\|Segment:S\|Host:Human |
| gb:EF990909\|Organism:Hantaanvirus CGHu3612\|Strain Name:CGHu3612\|Segment:S\|Host:Human |
| gb:EF990908\|Organism:Hantaanvirus CGHu3614\|Strain Name:CGHu3614\|Segment:S\|Host:Human |
| gb:MG923671\|Organism:Puumala orthohantavirus\|Strain Name:AISNE-02/Hu/FRA/2016.00467\|Segment:S\|Host:Human |
| gb:MG923604\|Organism:Puumala orthohantavirus\|Strain Name:ALFORTVILLE-94/Hu/FRA/2015.00456\|Segment:S\|Host:Human |
| gb:MG923608\|Organism:Puumala orthohantavirus\|Strain Name:ANGIREY-70/Hu/FRA/2015.00410\|Segment:S\|Host:Human |
| gb:MG923656\|Organism:Puumala orthohantavirus\|Strain Name:ANOR-59/Hu/FRA/2015.00422\|Segment:S\|Host:Human |
| gb:MG923652\|Organism:Puumala orthohantavirus\|Strain Name:ARBOIS-39/Hu/FRA/2014.00622\|Segment:S\|Host:Human |
| gb:MG923647\|Organism:Puumala orthohantavirus\|Strain Name:ATHIES-SOUS-LAON-02/Hu/FRA/2014.00135\|Segment:S\|Host:Human |
| gb:MG923665\|Organism:Puumala orthohantavirus\|Strain Name:AULNOYE-AYMERIES-59/Hu/FRA/2016.00325\|Segment:S\|Host:Human |
| gb:MG923605\|Organism:Puumala orthohantavirus\|Strain Name:BAR-LE-DUC-55/Hu/FRA/2012.00123\|Segment:S\|Host:Human |
| gb:MG923627\|Organism:Puumala orthohantavirus\|Strain Name:BOGNY-SUR-MEUSE-08/Hu/FRA/2015.00329\|Segment:S\|Host:Human |
| gb:MG923660\|Organism:Puumala orthohantavirus\|Strain Name:BOULZICOURT-08/Hu/FRA/2016.00182\|Segment:S\|Host:Human |
| gb:MG923618\|Organism:Puumala orthohantavirus\|Strain Name:BUIRONFOSSE-02/Hu/FRA/2014.00153\|Segment:S\|Host:Human |
| gb:MG923640\|Organism:Puumala orthohantavirus\|Strain Name:CESSIERES-02/Hu/FRA/2016.00353\|Segment:S\|Host:Human |
| gb:MG923623\|Organism:Puumala orthohantavirus\|Strain Name:CHAMBLY-60/Hu/FRA/2014.00540\|Segment:S\|Host:Human |
| gb:MG923600\|Organism:Puumala orthohantavirus\|Strain Name:CHAMPIGNY-SUR-MARNE-94/Hu/FRA/2014.00499\|Segment:S\|Host:Human |
| gb:MG923654\|Organism:Puumala orthohantavirus\|Strain Name:CHARLEVILLE-MEZIERES-08/Hu/FRA/2015.00402\|Segment:S\|Host:Human |
| gb:MG923611\|Organism:Puumala orthohantavirus\|Strain Name:CHEVROCHES-58/Hu/FRA/2012.00086\|Segment:S\|Host:Human |
| gb:MG923631\|Organism:Puumala orthohantavirus\|Strain Name:CILLY-02/Hu/FRA/2015.00657\|Segment:S\|Host:Human |
| gb:MG923606\|Organism:Puumala orthohantavirus\|Strain Name:COISERETTE-39/Hu/FRA/2012.00102\|Segment:S\|Host:Human |
| gb:MG923612\|Organism:Puumala orthohantavirus\|Strain Name:COLOMBEY-LES-BELLES-54/Hu/FRA/2012.00307\|Segment:S\|Host:Human |
| gb:MG923663\|Organism:Puumala orthohantavirus\|Strain Name:CORNY-MACHEROMENIL-08/Hu/FRA/2016.00295\|Segment:S\|Host:Human |
| gb:MG923641\|Organism:Puumala orthohantavirus\|Strain Name:COUSOLRE-59/Hu/FRA/2012.00057\|Segment:S\|Host:Human |
| gb:MG923655\|Organism:Puumala orthohantavirus\|Strain Name:DOUZY-08/Hu/FRA/2015.00419\|Segment:S\|Host:Human |
| gb:MG923644\|Organism:Puumala orthohantavirus\|Strain Name:ENGLANCOURT-02/Hu/FRA/2012.00349\|Segment:S\|Host:Human |
| gb:MG923624\|Organism:Puumala orthohantavirus\|Strain Name:ETEIGNIERES-08/Hu/FRA/2015.00019\|Segment:S\|Host:Human |
| gb:MG923626\|Organism:Puumala orthohantavirus\|Strain Name:FELLERING-68/Hu/FRA/2015.00185\|Segment:S\|Host:Human |
| gb:MG923649\|Organism:Puumala orthohantavirus\|Strain Name:FOURMIES-59/Hu/FRA/2014.00184\|Segment:S\|Host:Human |
| gb:MG923650\|Organism:Puumala orthohantavirus\|Strain Name:FOURMIES-59/Hu/FRA/2014.00233\|Segment:S\|Host:Human |
| gb:MG923622\|Organism:Puumala orthohantavirus\|Strain Name:FOURMIES-59/Hu/FRA/2014.00321\|Segment:S\|Host:Human |
| gb:MG923601\|Organism:Puumala orthohantavirus\|Strain Name:FOURMIES-59/Hu/FRA/2014.00598\|Segment:S\|Host:Human |
| gb:MG923651\|Organism:Puumala orthohantavirus\|Strain Name:FOURMIES-59/Hu/FRA/2014.00613\|Segment:S\|Host:Human |
| gb:MG923625\|Organism:Puumala orthohantavirus\|Strain Name:FOURMIES-59/Hu/FRA/2015.00045\|Segment:S\|Host:Human |
| gb:MG923666\|Organism:Puumala orthohantavirus\|Strain Name:FOURMIES-59/Hu/FRA/2016.00333\|Segment:S\|Host:Human |
| gb:MG923667\|Organism:Puumala orthohantavirus\|Strain Name:FOURMIES-59/Hu/FRA/2016.00345\|Segment:S\|Host:Human |
| gb:MG923669\|Organism:Puumala orthohantavirus\|Strain Name:FOURMIES-59/Hu/FRA/2016.00427\|Segment:S\|Host:Human |
| gb:MG923615\|Organism:Puumala orthohantavirus\|Strain Name:GIVET-08/Hu/FRA/2012.00638\|Segment:S\|Host:Human |
| gb:MG923614\|Organism:Puumala orthohantavirus\|Strain Name:GOUVIEUX-60/Hu/FRA/2012.00402\|Segment:S\|Host:Human |
| gb:MG923653\|Organism:Puumala orthohantavirus\|Strain Name:GREZY-SUR-ISERE-73/Hu/FRA/2015.00153\|Segment:S\|Host:Human |
| gb:MK496162\|Organism:Puumala orthohantavirus\|Strain Name:H46/Ufa\|Segment:S\|Host:Human |
| gb:MG923668\|Organism:Puumala orthohantavirus\|Strain Name:HIRSON-02/Hu/FRA/2016.00357\|Segment:S\|Host:Human |
| gb:MG923633\|Organism:Puumala orthohantavirus\|Strain Name:JALLANGES-21/Hu/FRA/2016.00275\|Segment:S\|Host:Human |
| gb:MG923635\|Organism:Puumala orthohantavirus\|Strain Name:LA-NEUVILLE-SUR-RESSONS-60/Hu/FRA/2016.00293\|Segment:S\|Host:Human |
| gb:MG923645\|Organism:Puumala orthohantavirus\|Strain Name:LA-PESSE-39/Hu/FRA/2012.00536\|Segment:S\|Host:Human |
| gb:MG923609\|Organism:Puumala orthohantavirus\|Strain Name:LANISCOURT-02/Hu/FRA/2012.00061\|Segment:S\|Host:Human |
| gb:MG923636\|Organism:Puumala orthohantavirus\|Strain Name:LAON-02/Hu/FRA/2016.00311\|Segment:S\|Host:Human |
| gb:MG923639\|Organism:Puumala orthohantavirus\|Strain Name:LAON-02/Hu/FRA/2016.00326\|Segment:S\|Host:Human |
| gb:MG923670\|Organism:Puumala orthohantavirus\|Strain Name:LAON-02/Hu/FRA/2016.00452\|Segment:S\|Host:Human |
| gb:MG923607\|Organism:Puumala orthohantavirus\|Strain Name:LE-MOUTARET-38/Hu/FRA/2014.00120\|Segment:S\|Host:Human |
| gb:MG923621\|Organism:Puumala orthohantavirus\|Strain Name:LILLE-59/Hu/FRA/2014.00276\|Segment:S\|Host:Human |
| gb:MG923628\|Organism:Puumala orthohantavirus\|Strain Name:MONTCORNET-02/Hu/FRA/2015.00430\|Segment:S\|Host:Human |
| gb:MG923630\|Organism:Puumala orthohantavirus\|Strain Name:MONTHERME-08/Hu/FRA/2015.00526\|Segment:S\|Host:Human |
| gb:MG923634\|Organism:Puumala orthohantavirus\|Strain Name:MORBECQUE-59/Hu/FRA/2016.00282\|Segment:S\|Host:Human |
| gb:MG923610\|Organism:Puumala orthohantavirus\|Strain Name:MOUTHE-25/Hu/FRA/2012.00301\|Segment:S\|Host:Human |
| gb:MK496159\|Organism:Puumala orthohantavirus\|Strain Name:P-360\|Segment:S\|Host:Human |
| gb:MG923672\|Organism:Puumala orthohantavirus\|Strain Name:PREMONTRE-02/Hu/FRA/2016.00469\|Segment:S\|Host:Human |
| gb:MG923661\|Organism:Puumala orthohantavirus\|Strain Name:PRESLES-ET-THIERNY-02/Hu/FRA/2016.00268\|Segment:S\|Host:Human |
| gb:MH251331\|Organism:Puumala orthohantavirus\|Strain Name:PUU-TKD\|Segment:S\|Host:Human |
| gb:JN831950\|Organism:Puumala orthohantavirus\|Strain Name:PUUV/Pieksamaki/human_kidney/2008\|Segment:S\|Host:Human |
| gb:JN831947\|Organism:Puumala orthohantavirus\|Strain Name:PUUV/Pieksamaki/human_lung/2008\|Segment:S\|Host:Human |
| gb:MG923643\|Organism:Puumala orthohantavirus\|Strain Name:REIMS-51/Hu/FRA/2012.00278\|Segment:S\|Host:Human |
| gb:MG923674\|Organism:Puumala orthohantavirus\|Strain Name:REIMS-51/Hu/FRA/2015.00665\|Segment:S\|Host:Human |
| gb:MG923658\|Organism:Puumala orthohantavirus\|Strain Name:REMILLY-AILLICOURT-08/Hu/FRA/2015.00498\|Segment:S\|Host:Human |
| gb:MG923629\|Organism:Puumala orthohantavirus\|Strain Name:REVIGNY-SUR-ORNAIN-55/Hu/FRA/2015.00457\|Segment:S\|Host:Human |
| gb:MG923598\|Organism:Puumala orthohantavirus\|Strain Name:RIOZ-70/Hu/FRA/2015.00567\|Segment:S\|Host:Human |
| gb:MG923673\|Organism:Puumala orthohantavirus\|Strain Name:ROCROI-08/Hu/FRA/2012.00018\|Segment:S\|Host:Human |
| gb:MG923638\|Organism:Puumala orthohantavirus\|Strain Name:RONCHAMP-70/Hu/FRA/2015.00504\|Segment:S\|Host:Human |
| gb:MG923613\|Organism:Puumala orthohantavirus\|Strain Name:SAINT-CLAUDE-39/Hu/FRA/2012.00396\|Segment:S\|Host:Human |
| gb:MG923646\|Organism:Puumala orthohantavirus\|Strain Name:SAINT-MICHEL-02/Hu/FRA/2014.00097\|Segment:S\|Host:Human |
| gb:MG923619\|Organism:Puumala orthohantavirus\|Strain Name:SAINT-SAULVE-59/Hu/FRA/2014.00171\|Segment:S\|Host:Human |
| gb:MG923637\|Organism:Puumala orthohantavirus\|Strain Name:SAINT-VIT-25/Hu/FRA/2016.00320\|Segment:S\|Host:Human |
| gb:MG923603\|Organism:Puumala orthohantavirus\|Strain Name:SAINTE-MENEHOULD-51/Hu/FRA/2012.00025\|Segment:S\|Host:Human |
| gb:MG923659\|Organism:Puumala orthohantavirus\|Strain Name:SAULES-25/Hu/FRA/2014.00637\|Segment:S\|Host:Human |
| gb:MG923617\|Organism:Puumala orthohantavirus\|Strain Name:SECHEVAL-08/Hu/FRA/2014.00053\|Segment:S\|Host:Human |
| gb:MG923657\|Organism:Puumala orthohantavirus\|Strain Name:SEDAN-08/Hu/FRA/2015.00488\|Segment:S\|Host:Human |
| gb:MG923642\|Organism:Puumala orthohantavirus\|Strain Name:SIGNY-LE-PETIT-08/Hu/FRA/2014.00488\|Segment:S\|Host:Human |
| gb:MG923648\|Organism:Puumala orthohantavirus\|Strain Name:ST-ERME-OUTRE-ET-RAMECOURT-02/Hu/FRA/2014.00174\|Segment:S\|Host:Human |
| gb:MG923664\|Organism:Puumala orthohantavirus\|Strain Name:THIN-LE-MOUTIER-08/Hu/FRA/2016.00310\|Segment:S\|Host:Human |
| gb:MG923620\|Organism:Puumala orthohantavirus\|Strain Name:TREMBLOIS-LES-ROCROI-08/Hu/FRA/2014.00209\|Segment:S\|Host:Human |
| gb:MG923662\|Organism:Puumala orthohantavirus\|Strain Name:TRUCY-02/Hu/FRA/2016.00286\|Segment:S\|Host:Human |
| gb:MG923616\|Organism:Puumala orthohantavirus\|Strain Name:VENDIN-LES-BETHUNE-62/Hu/FRA/2013.00250\|Segment:S\|Host:Human |
| gb:MG923599\|Organism:Puumala orthohantavirus\|Strain Name:VIC-SUR-AISNE-02/Hu/FRA/2015.00660\|Segment:S\|Host:Human |
| gb:MG923632\|Organism:Puumala orthohantavirus\|Strain Name:VIREUX-MOLHAIN-08/Hu/FRA/2016.00239\|Segment:S\|Host:Human |
| gb:MG923602\|Organism:Puumala orthohantavirus\|Strain Name:VRIGNE-MEUSE-08/Hu/FRA/2015.00328\|Segment:S\|Host:Human |
| gb:GQ279395\|Organism:Seoul orthohantavirus\|Strain Name:CUI\|Segment:S\|Host:Human |
| gb:KX064275\|Organism:Seoul orthohantavirus\|Strain Name:ERIZE-ST-DIZIER/Hu/FRA/2014/2014.00479\|Segment:S\|Host:Human |
| gb:MF149954\|Organism:Seoul orthohantavirus\|Strain Name:Hu02-258/NGS\|Segment:S\|Subtype:Seoul\|Host:Human |
| gb:MF149955\|Organism:Seoul orthohantavirus\|Strain Name:Hu02-294/NGS\|Segment:S\|Subtype:Seoul\|Host:Human |
| gb:MF149956\|Organism:Seoul orthohantavirus\|Strain Name:Hu02-529/NGS\|Segment:S\|Subtype:Seoul\|Host:Human |
| gb:GQ279390\|Organism:Seoul orthohantavirus\|Strain Name:HuBJ15\|Segment:S\|Host:Human |
| gb:GQ279380\|Organism:Seoul orthohantavirus\|Strain Name:HuBJ16\|Segment:S\|Host:Human |
| gb:GQ279389\|Organism:Seoul orthohantavirus\|Strain Name:HuBJ19\|Segment:S\|Host:Human |
| gb:GQ279394\|Organism:Seoul orthohantavirus\|Strain Name:HuBJ20\|Segment:S\|Host:Human |
| gb:GQ279379\|Organism:Seoul orthohantavirus\|Strain Name:HuBJ22\|Segment:S\|Host:Human |
| gb:GQ279391\|Organism:Seoul orthohantavirus\|Strain Name:HuBJ3\|Segment:S\|Host:Human |
| gb:GQ279381\|Organism:Seoul orthohantavirus\|Strain Name:HuBJ7\|Segment:S\|Host:Human |
| gb:GQ279384\|Organism:Seoul orthohantavirus\|Strain Name:HuBJ9\|Segment:S\|Host:Human |
| gb:KC902522\|Organism:Seoul orthohantavirus\|Strain Name:REPLONGES/Hu/FRA/2012/12-0882\|Segment:S\|Host:Human |
| gb:KX064270\|Organism:Seoul orthohantavirus\|Strain Name:TURCKHEIM/Hu/FRA/2016/2016.00044\|Segment:S\|Host:Human |
| gb:KT946591\|Organism:Tula orthohantavirus\|Strain Name:CHEVRU/Hu/FRA/2015/15.00453\|Segment:S\|Host:Human |

**Figure 9.**
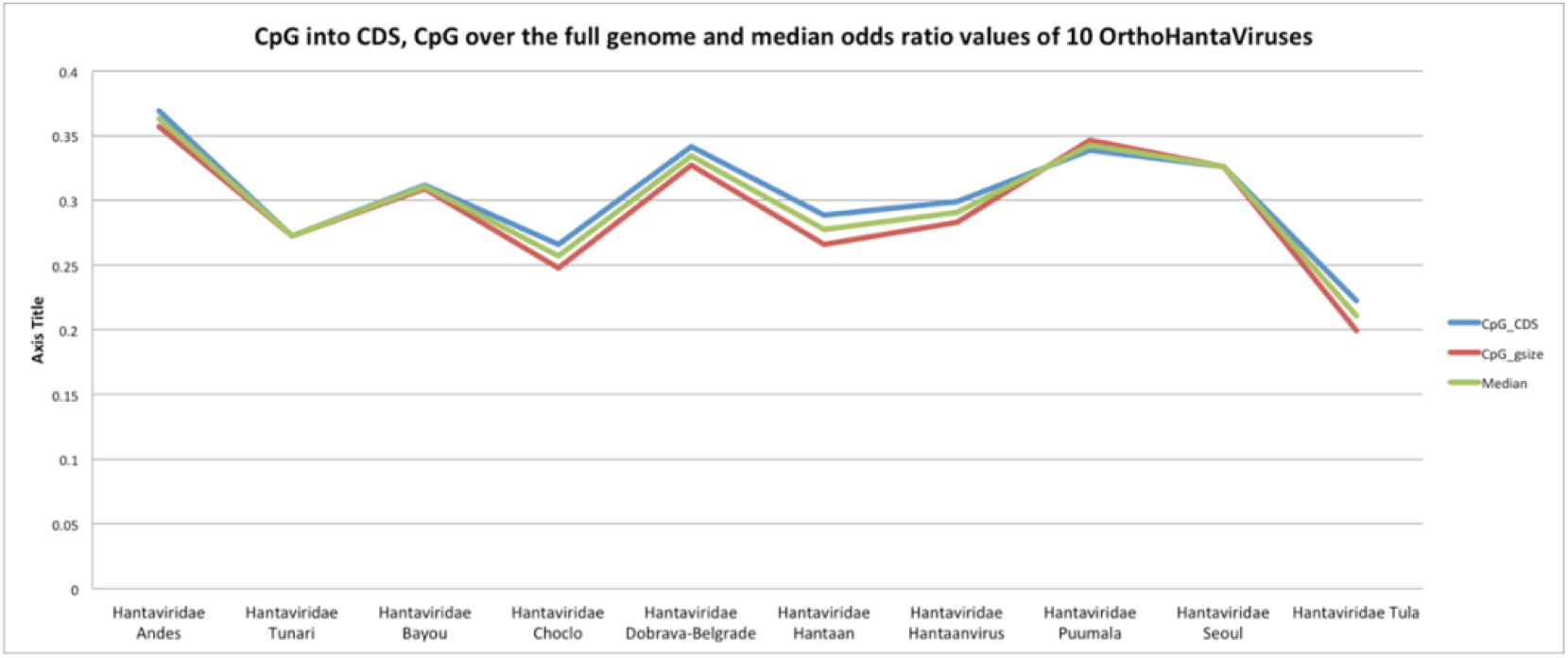
The Andes Hantaviridae shows the highest values in all the three cases (CpG odds ratio into CDS, CpG odds ratio from full genome and Median values)

### Optimal number of clusters for Hantaviruses

As mentioned in the Method section, we used the Elbow curve, Silhouette score, Gap statistic, and clustree discovery to investigate the optimal number of clusters (k).

The Elbow curve method looks at the total within-cluster sum of squares (WSS) as a function of the number of clusters. We considered as an indicator of the appropriate k value the location of a knee in the plot. The Elbow method suggested k=4 as the optimal partitioning. The Silhouette score method measures the quality of clustering and determines how well each point lies within its cluster, and in our case, it suggested k=2. The optimal k is the one that maximizes the gap statistic. Approaching the problem by the GAP statistical method, it proposed only 1 cluster (which is a useless clustering). Figure 10 reports all three results.

**Figure 10.**
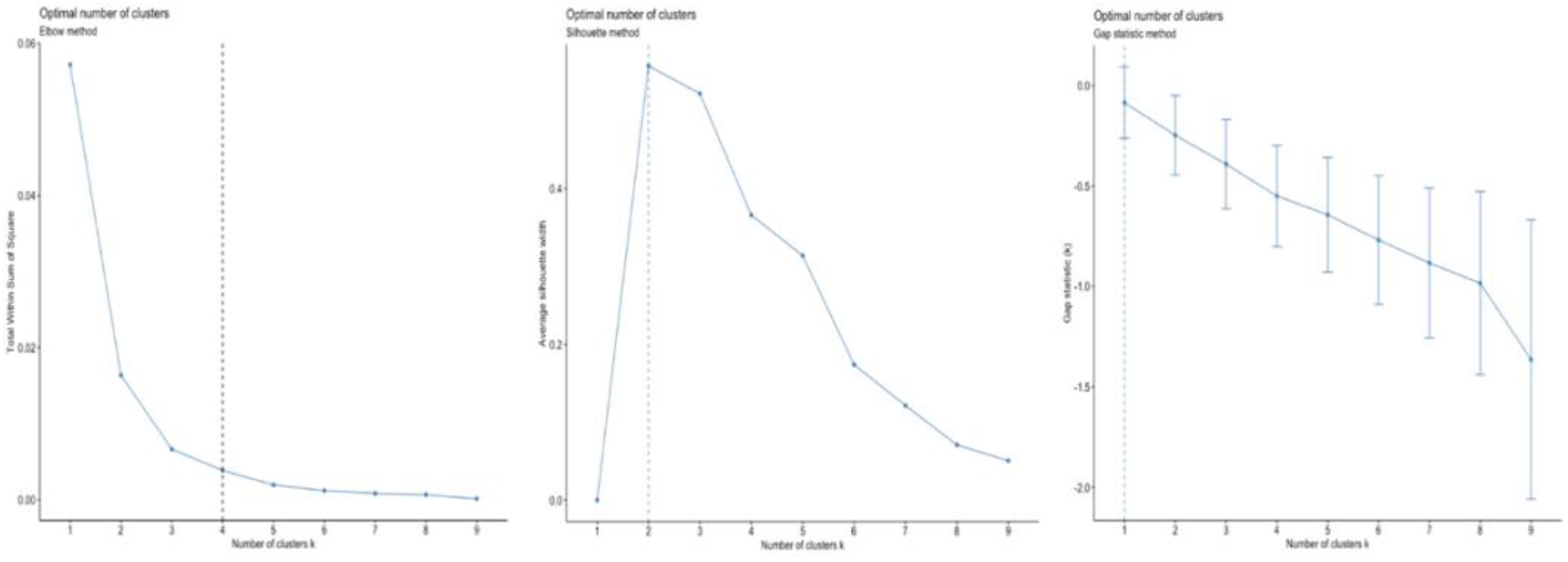
Optimal number of clusters according to Elbow, Silhouette and GAP methods

All three approaches suggested a different number of clusters, so we have used the discovery clustree approach to consider how samples change groupings as the number of clusters increases. This approach is useful for showing which clusters are distinct and unstable, exploring possible choices.

In Figure 11, the size of each node corresponds to the number of samples in each cluster. It also colors the arrows according to the number of samples each group receives. In this graph, passing from *k=2 to k=3*, several viruses are reassigned from the lookers-left cluster to the third cluster on the right. Moving from *k=4 to k=5*, two nodes present multiple incoming edges, indicating data over-clustering. Results show enough reasons to set *k=4* as the optimal number of clusters for our dataset.

**Figure 11.**
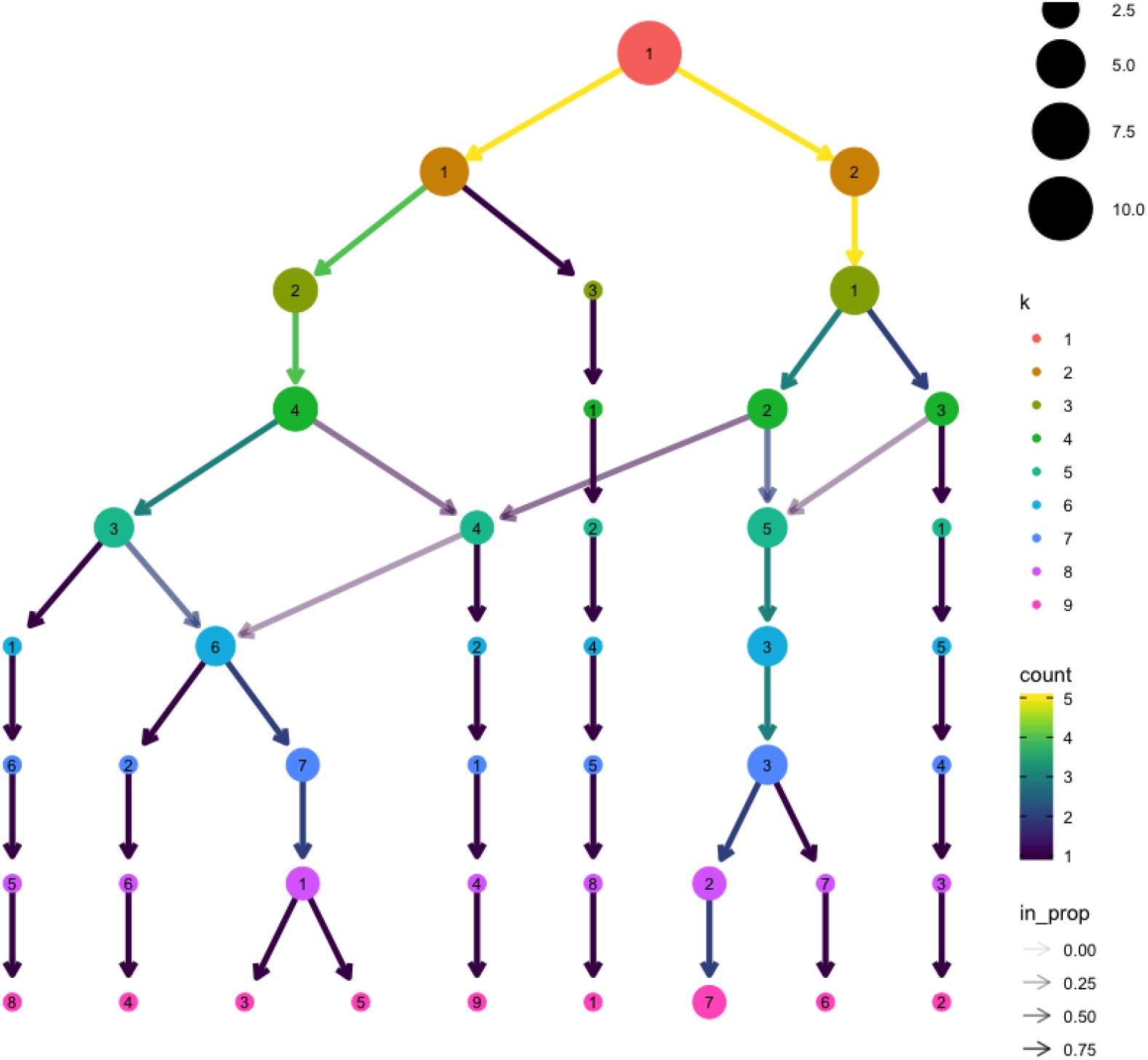
Cluster tree representation

### K-means, DBSCAN and HCA vs Hantavirus

We executed the K-means, DBSCAN, and HCA algorithms to find the groups of Hantaviruses more similar according to the CpG odds ratio, both from the full genome and the CDS regions, and concerning their median values from the group of small genomic segments. We focused attention on the Andes Hantavirus, being the unique hantavirus able to pass from human to human. K-mean algorithm showed the Andes H. as an element of the 4th cluster with the Puumala H. but showing a relevant distance from it (Figure 12). DBSCAN algorithm showed four groups of viruses, even if it does not well remark the distance between them (Figure 13). HCA agglomerative and divisive reported the same dendrogram, showing that Andes H. is a “borderline” virus as the Tula H., even if belonging to two different clusters (Figure 14). Representing the clustering obtained by the hierarchical methods, we got again evidence that Andes H. looks like an isolated cluster (as the Tula H.), suggesting the critical contrast with the other viruses from the same *Hantaviridae* family (Figure 15)

**Figure 12.**
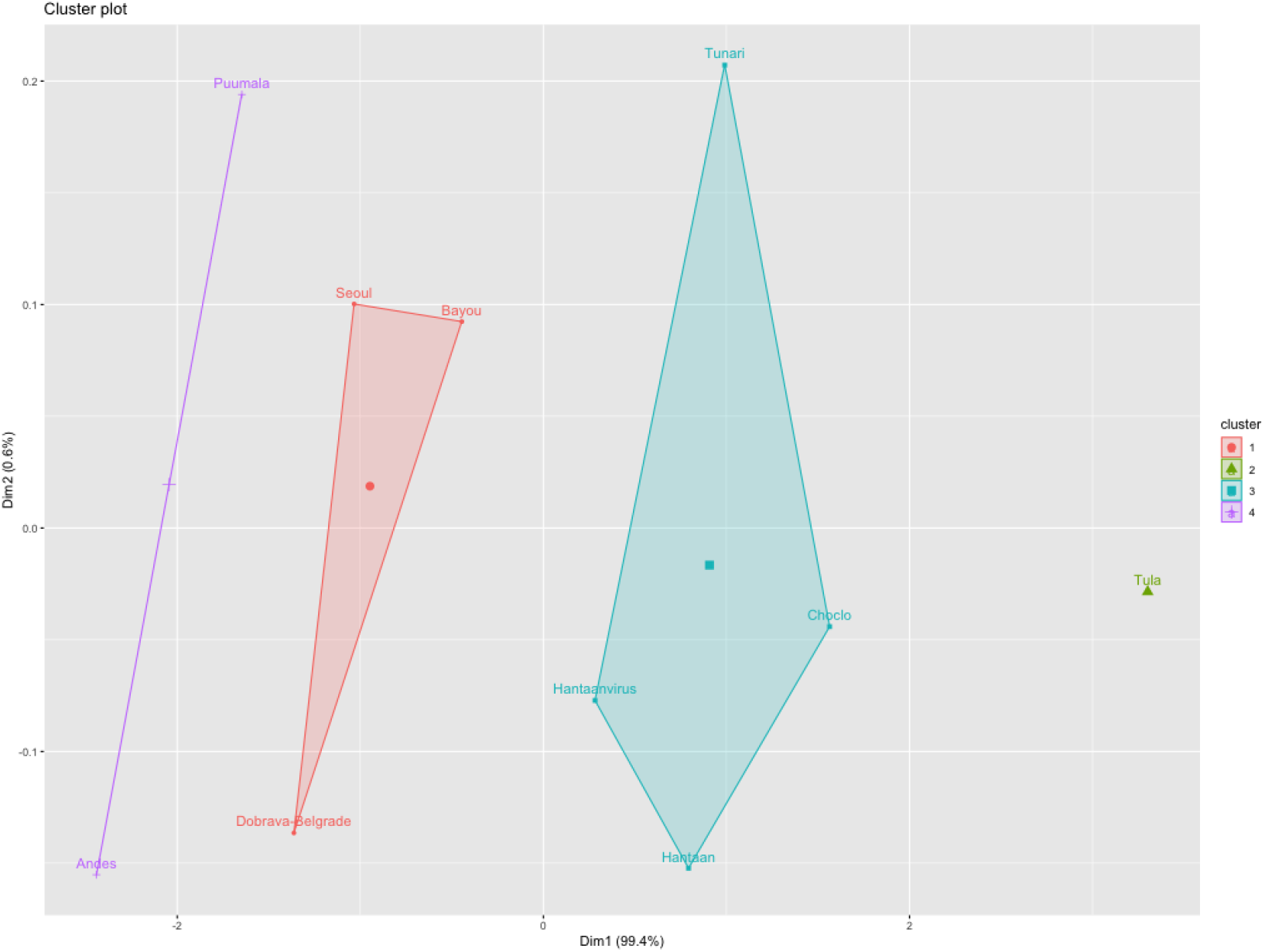
K-means with k=4

**Figure 13.**
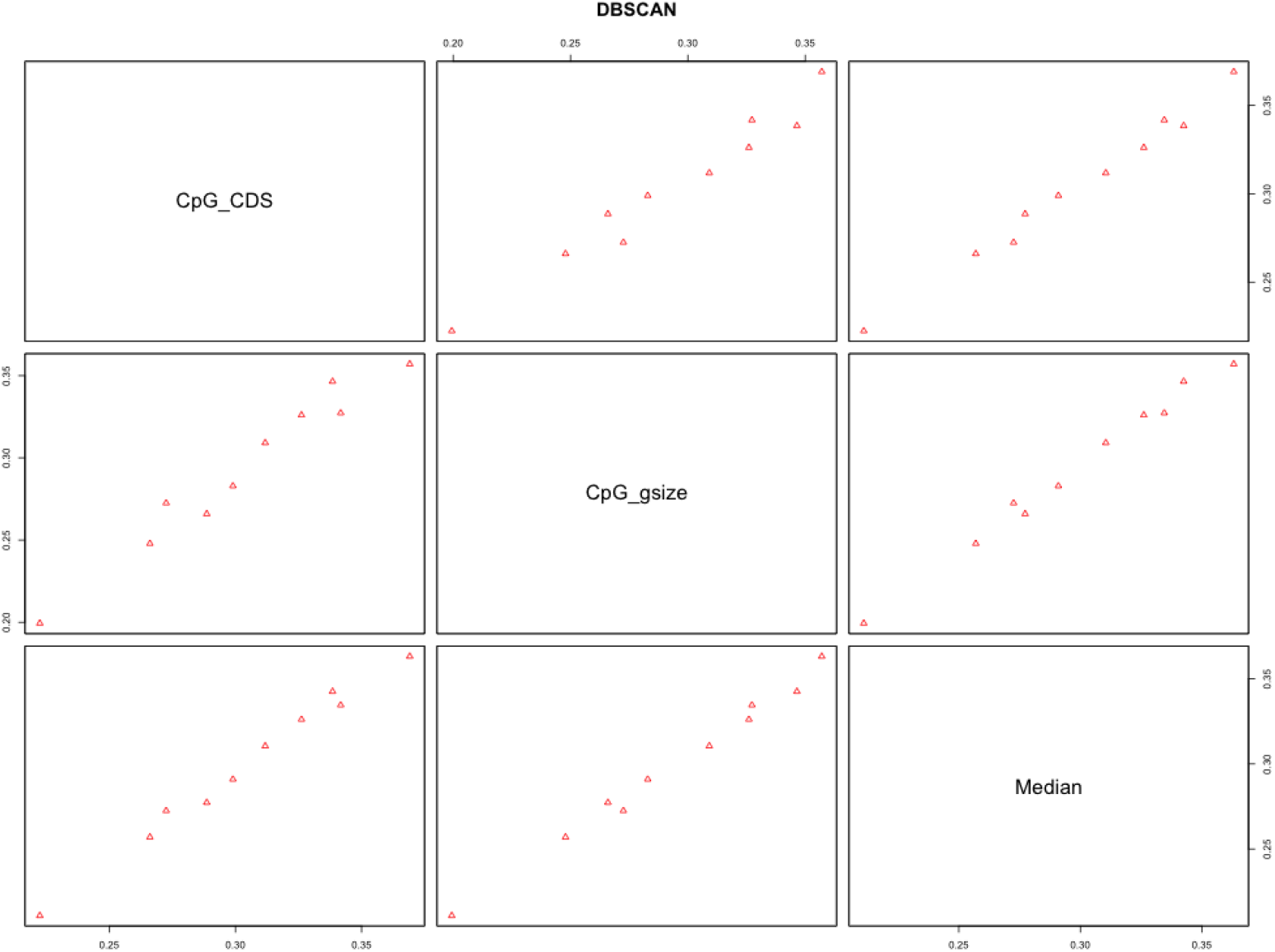
DBSCAN and four groups of viruses

**Figure 14.**
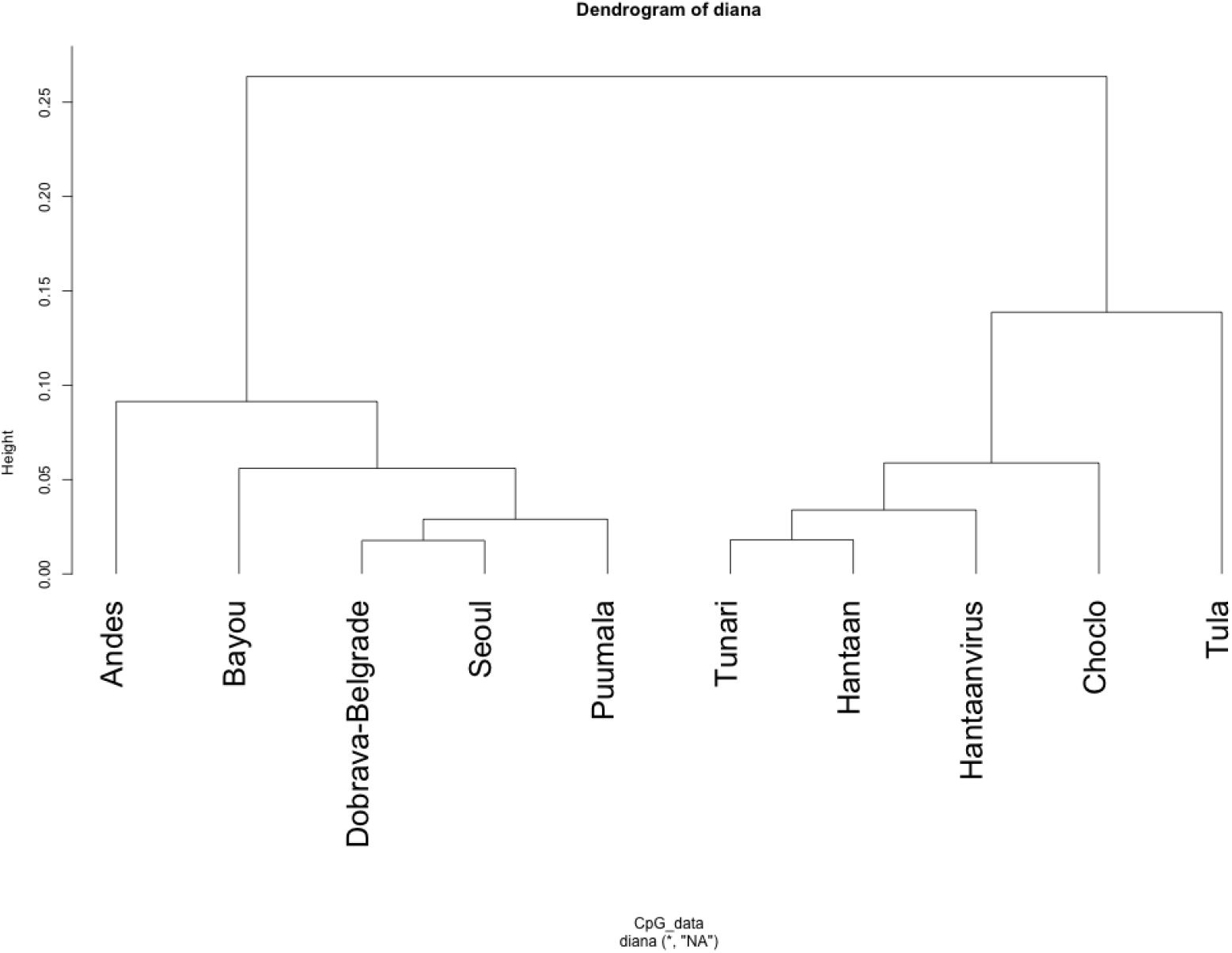
HCA divisive (AGNES)

**Figure 15.**
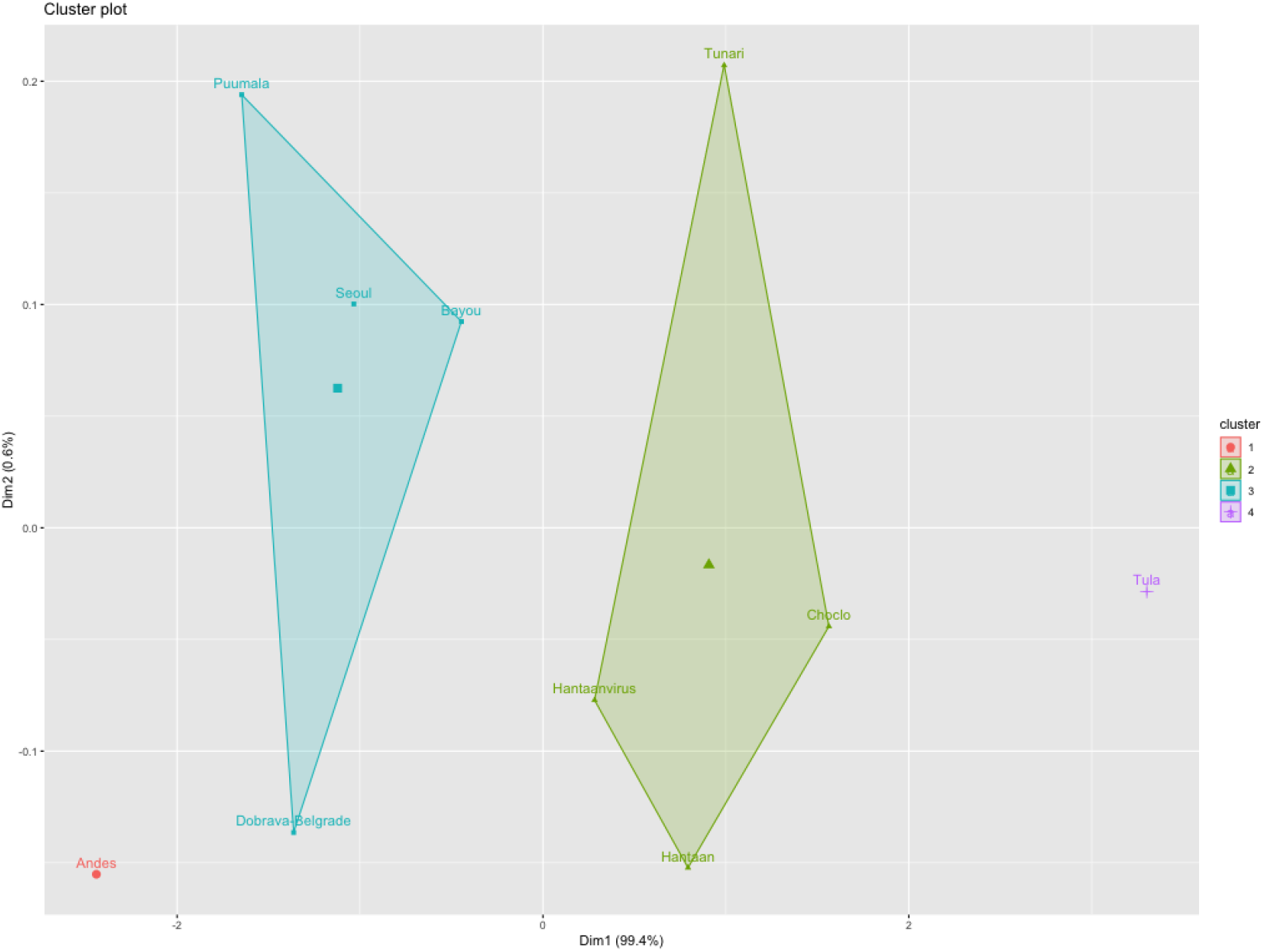
HCA clustering

**Figure 16.**
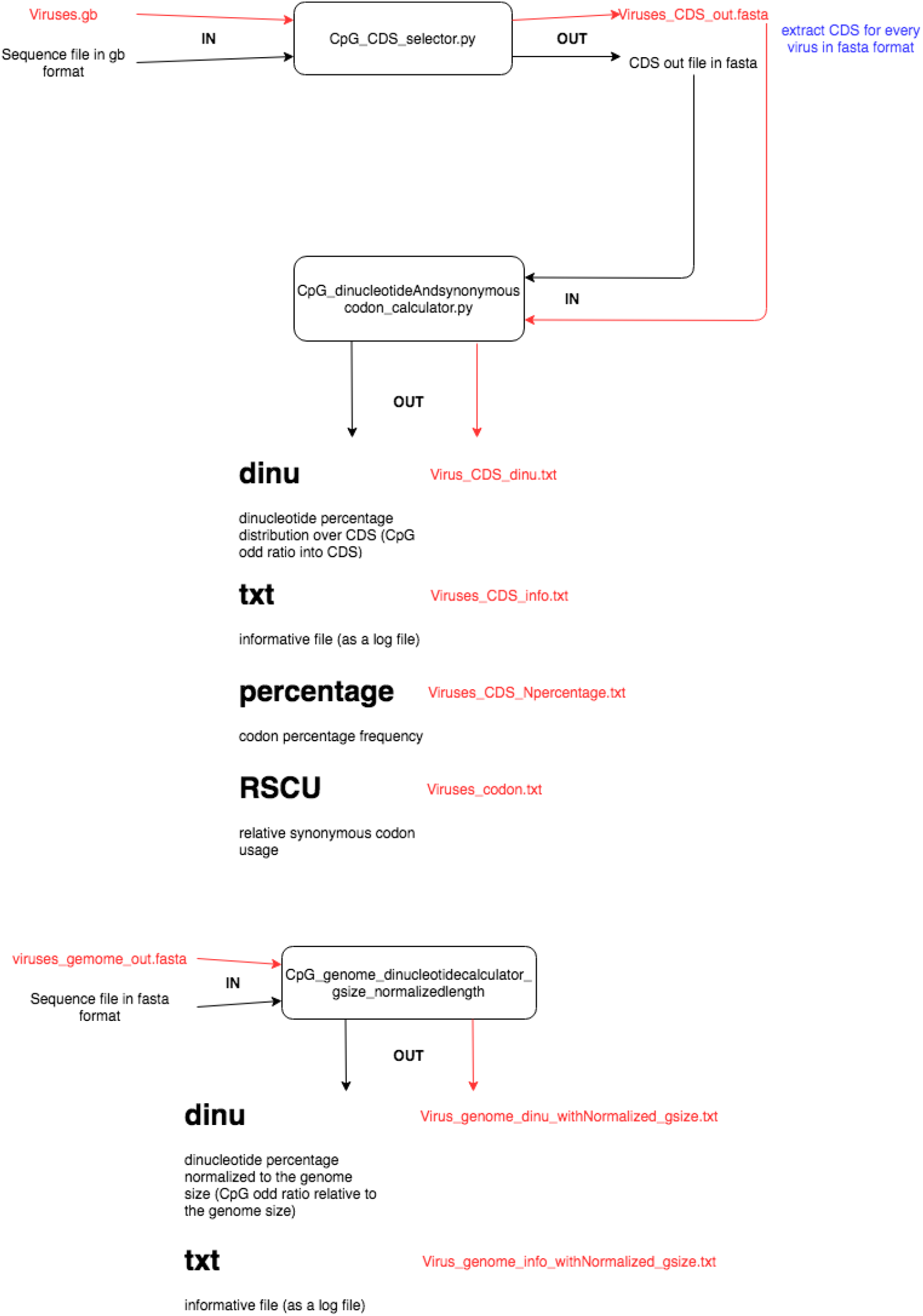
Flowchart of executed steps to calculate the CpG odds ratio

**Figure 17.**
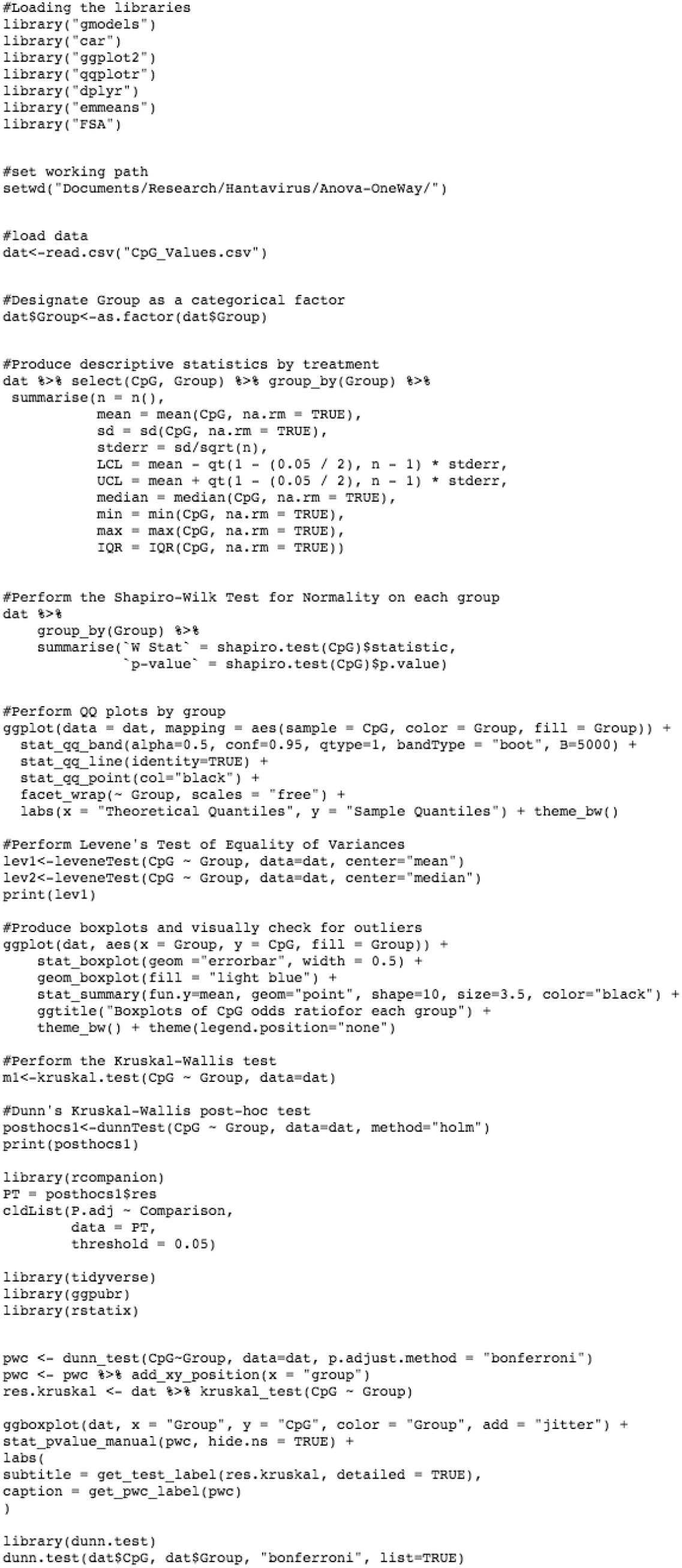
Script to conduct ANOVA analysis in R

**Figure 18.**
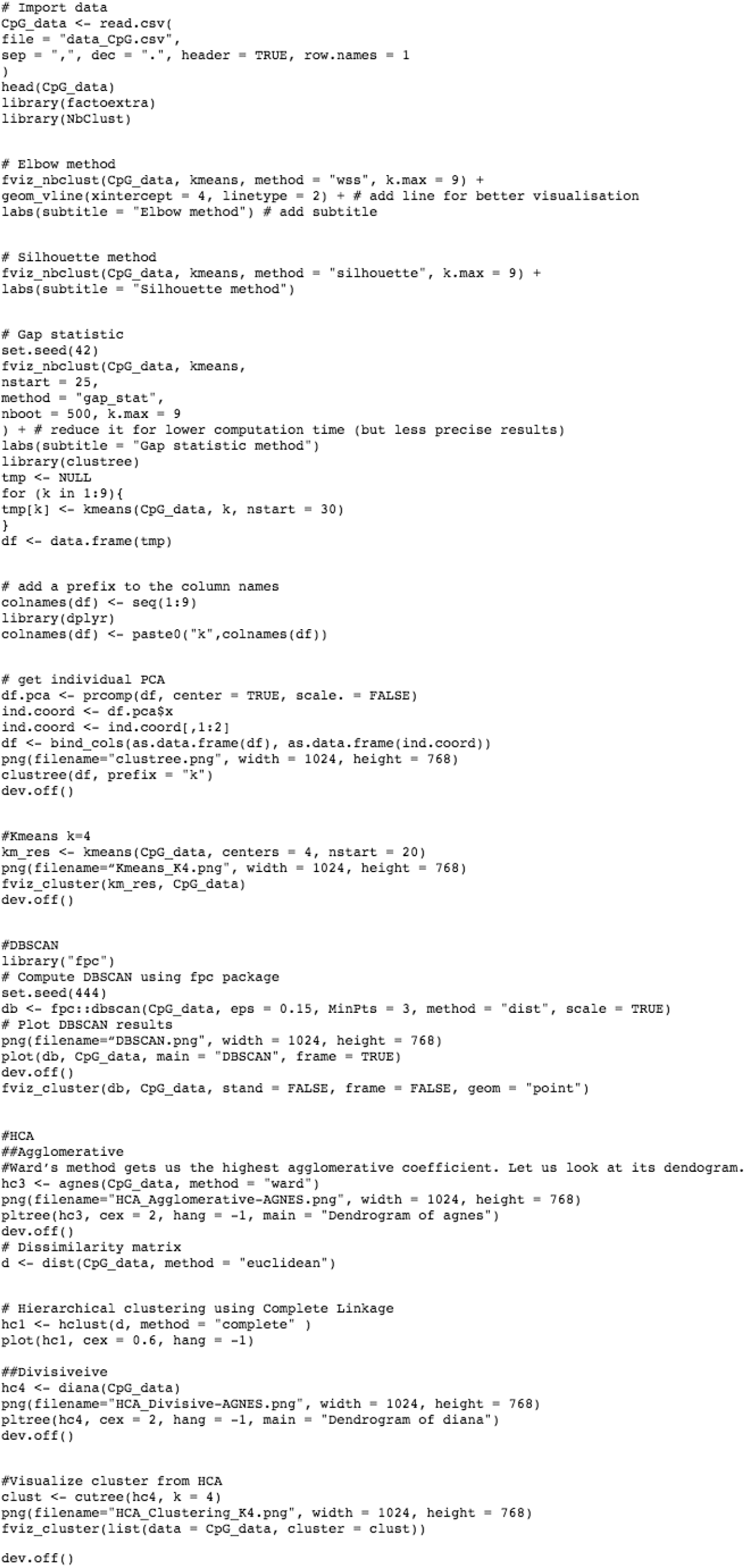
Script to conduct the unsupervised clustering in R

## Discussion and Conclusions

In the current study, we analyzed the *Orthohantaviridae* family from the CpG odds ratio point of view. As the first result, we proved the statistical difference between the three groups of segmented genomes. We have identified the small genomic segments group as the more informative, giving us the chance to reduce the research space. To avoid the influence of the CpG odds ratio from the non-coding regions, we first calculated the CpG odds ratio into the coding regions for the small segments of all viruses. As a result, we obtained the confirmation that the CpG frequency is the lowest compared to the other dinucleotides and that the Andes Hantavirus showed its highest CpG odds ratio in CDS.

Secondly, we considered the correlation coefficient between the CpG odds ratio and the CpG odds ratio of the coding regions. We performed the calculation from the small genomic groups of all the viruses, keeping in mind that a positive correlation implies a more significant CpG odds ratio from the small genomic segment group.

The correlation analysis between the CpG odds ratio from the full size of the segmented small genome and the CDS regions resulted in a positive index. This result underlines the possible function of the CpG islands inside of the coding regions. Comparing the CpG over the full genome, the CpG over the CDS and the median values over the ten viruses suggested a stronger concentration of the CpG islands both along the full-size genome and the CDS regions into the Andes virus. Using both the CpG odds ratio measurements (based on the CDS and full genome size) from the group of small genomic segments as features, the unsupervised clustering analysis identified four different sub-groups inside of the Orthohantaviridae family. The unsupervised clustering corroborated the evidence that the Andes Hantavirus (similar, in some way, to Tula H.) exhibits a peculiar CpG odds ratio distribution, perhaps linked to its unique prerogative to pass from human-to-human. Previous research already pointed out the huge variations of CpG bias in RNA viruses and brought out the observed under-representation of CpG in RNA viruses as not caused by the biased CpG usage in the non-coding regions but determined by the coding regions [12, 13]. The current study suggests that the prerogative of Andes H. to be transmitted from human to human could be linked to its distribution of CpG dinucleotides. The research pointed out that in any case, the frequency of CpG islands in the Andes H. virus is such as to be identified as a cluster in its own right. In the case of Tula orthohantavirus, infections being rarely found in humans [23–25] and even if (at the moment) there is no evidence to suggest diversification of this virus from the rest of the family, it is questionable whether this similarity suggests a potential anthroponotic capacity in this virus. We can certainly assert that even in its case the distribution of CpG dinucleotides suggests greater attention. As a possible step forward in the research carried out, surely the use of further features related to the distribution of CpG dinucleotides as a relationship index with the CpG distribution of the host or with the distribution of the CpG islands in the regions internal to the codons and between the codons could provide more detailed clustering results. The research carried out has already presented many important results, such as the significant statistical difference between the distributions of CpG dinucleotides in the different genomic segments (S, M, and L), the identification of numerical indices useful for the application of unsupervised clustering algorithms, and the identification of subgroups within the family of orthohantaviruses. These subgroups included Andes H. and Tula H. as cases worthy of particular attention, especially in reference to the Andes H. whose peculiar *anthroponicity* is particularly dangerous for humans.

